# Fragment-based screening identifies molecules targeting the substrate-binding ankyrin repeat domains of tankyrase

**DOI:** 10.1101/567446

**Authors:** Katie Pollock, Manjuan Liu, Mariola Zaleska, Mark Pfuhl, Ian Collins, Sebastian Guettler

## Abstract

The PARP enzyme and scaffolding protein tankyrase (TNKS, TNKS2) uses its ankyrin repeat clusters (ARCs) to bind a wide range of proteins and thereby controls diverse cellular functions. A number of these are implicated in cancer-relevant processes, including Wnt/β-catenin signaling and telomere maintenance. The ARCs recognise a conserved tankyrase-binding peptide motif (TBM). All currently available tankyrase inhibitors target the catalytic domain and inhibit tankyrase’s poly(ADP-ribosyl)ation function. However, there is emerging evidence that catalysis-independent “scaffolding” mechanisms contribute to tankyrase function. Here we report a fragment-based screening program against tankyrase ARC domains, using a combination of biophysical assays, including differential scanning fluorimetry (DSF) and nuclear magnetic resonance (NMR). We identify fragment molecules that will serve as starting points for the development of tankyrase substrate binding antagonists. Such compounds will enable probing the scaffolding functions of tankyrase, and may, in the future, provide potential alternative therapeutic approaches to inhibiting tankyrase activity in cancer and other conditions.

## Introduction

Tankyrase enzymes (TNKS/ARTD5 and TNKS2/ARTD6; simply referred to as ‘tankyrase’ from here on; Figure 1A) are poly(ADP-ribose)polymerases (PARPs) in the Diphtheria-toxin-like ADP-ribosyltransferase (ARTD) family ^1,2^. PARPs catalyse the processive addition of poly(ADP-ribose) (PAR) onto substrate proteins, which can either directly regulate acceptor protein function or serve as docking sites for PAR-binding proteins that mediate downstream signaling ^3^. Given the diversity of tankyrase binders and substrates, tankyrase impinges on a wide range of cellular functions ^2,4-6^. These include Wnt/β-catenin signaling ^7-10^, telomerase-dependent telomere lengthening ^11,12^, sister telomere resolution during mitosis ^13,14^, the control of glucose homeostasis ^15-17^, mitotic spindle assembly ^18,19^, DNA repair ^20,21^, and the regulation of the tumor-suppressive Hippo signaling pathway ^22-24^. Silencing of tankyrase elicits synthetic lethality in BRCA1/2-deficient cancer cells ^25^. Given these links of tankyrase to disease-relevant processes, tankyrase has gained attention as a potential therapeutic target ^2,26^.

**Figure 1.**
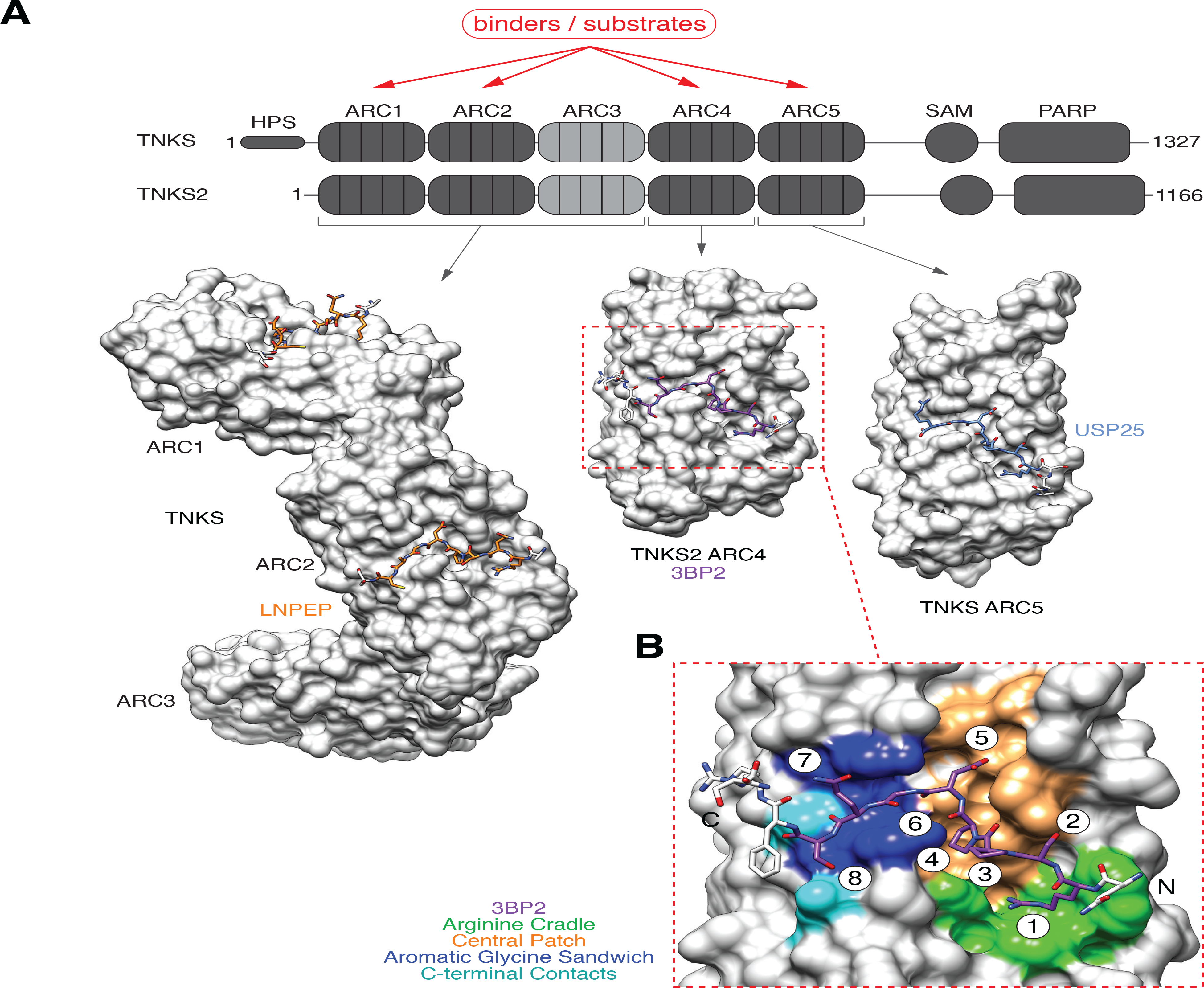
**(A)** Domain organisation of human tankyrase enzymes. Two tankyrase paralogs (TNKS, TNKS2) share an overall sequence identity of 82% (83% across ARCs, 74% across SAM domains, 94% across PARP domains). The ARCs comprise the substrate/protein recognition domain. Several examples of crystal structures of human ARCs bound to tankyrase-binding motif (TBM) peptides are shown: TNKS ARC1-3 bound to TBM peptide from LNPEP ^48^ (PDB code 5JHQ), TNKS2 ARC4 bound to TBM peptide from 3BP2 ^4^ (3TWR), TNKS ARC5 bound to TBM peptide from USP25 ^63^ (5GP7). **(B)** Details of the interaction of TNKS2 ARC4 with a 3BP2 TBM peptide ^4^ (3TWR). Four TBM peptide-binding hotspots are shown: (1) the “arginine cradle” (green), (2) “central patch” (orange), (3) “aromatic glycine sandwich” (blue), and (4) “C-terminal contacts” (cyan). TBM octapeptide amino acid positions are numbered.

Many mechanistic aspects of tankyrase function have been revealed by studying its role in Wnt/β-catenin signaling ^10^. Tankyrase promotes Wnt/β-catenin signaling by PARylating AXIN (axis inhibition protein 1/2) ^7^, a central component of the β-catenin destruction complex, which induces the degradation of the transcriptional co-activator β-catenin under low-Wnt conditions ^27^. AXIN PARylation either induces its PAR-dependent ubiquitination and degradation ^7,28-30^, or promotes the Wnt-induced transformation of the destruction complex into a signalosome complex incapable of initiating β-catenin degradation ^8,31^. Tankyrase thus sensitises cells to incoming Wnt signals ^32,33^. The Wnt/β-catenin pathway is dysregulated in the vast majority of colorectal cancers ^34^. Inhibiting tankyrase has been explored as a strategy to re-tune oncogenically dysregulated Wnt/β-catenin signaling in cancers with mutations in the tumor suppressor and destruction complex component APC (adenomatous polyposis coli) ^10,35-40^. Whilst tankyrase inhibitors can suppress tumor cell growth, *in-vivo* studies have also pointed to different degrees of tankyrase-inhibitor-induced intestinal toxicity in mice ^35,39,41^. The precise molecular mechanisms by which tankyrase controls Wnt/β-catenin signaling, how tankyrase inhibition can restore oncogenically dysregulated signaling, and the basis of tankyrase inhibitor toxicity are incompletely charted. This warrants the development of different chemical probes to modulate tankyrase function.

To date, drug discovery efforts on tankyrase have focused on inhibiting the catalytic PARP domain ^2,10,42,43^. Catalytic inhibition of tankyrase has complex consequences. As well as inhibiting substrate PARylation, catalytic inhibitors prevent tankyrase auto-PARylation and therefore subsequent PAR-dependent ubiquitination and degradation of tankyrase itself ^7^. Consequently, tankyrase inhibition typically leads not only to the accumulation of many of its substrates but also of tankyrase itself ^6,7,35,37,40^. Moreover, catalysis-independent functions of tankyrase are emerging, and these may be accentuated when tankyrase and its substrates accumulate upon tankyrase catalytic inhibition. Surprisingly, we observed that tankyrase can promote Wnt/β-catenin signaling independently of its catalytic PARP activity, at least when tankyrase levels are high ^9^. Under these conditions, tankyrase catalytic inhibitors only incompletely block tankyrase-driven β-catenin-dependent transcription, pointing to both catalytic and non-catalytic (scaffolding) functions of tankyrase. Tankyrase scaffolding functions depend on tankyrase’s substrate-binding ankyrin repeat clusters (ARCs) and the polymerization function of its sterile alpha motif (SAM) domain (see Figure 1A) ^9^. Tankyrase auto-PARylation has been proposed to limit tankyrase polymerization ^44^; tankyrase catalytic inhibition may therefore induce its hyperpolymerisation, which may further promote scaffolding functions. Scaffolding functions of tankyrase likely extend beyond Wnt/β-catenin signalling: not all tankyrase binders are also PARylated, and non-catalytic roles of tankyrase in other processes have been proposed ^4,5,45,46^.

Unraveling the complexity of tankyrase’s catalytic vs. non-catalytic functions will require novel tool compounds that block tankyrase-dependent scaffolding. Here, we identify and characterize small molecule fragments that bind to the tankyrase ARC domains, as a first step towards the discovery of compounds capable of blocking the interaction of tankyrase binders and substrates with the ARC domains of tankyrase.

Tankyrase contains five N-terminal ankyrin repeat clusters (ARCs), four of which, namely ARCs 1, 2, 4 and 5, can recruit binders and substrates ^4,47,48^ (Figure 1A). ARCs bind conserved but degenerate six-to eight-amino-acid peptide motifs, termed the tankyrase-binding motif (TBM, consensus R-X-X-[small hydrophobic or G]-[D/E/I/P]-G-[no P]-[D/E]) ^4,49^. Depending on the binding partner, ARCs can be functionally redundant, at least at the level of substrate recruitment ^4^, or collaborate in a combinatorial fashion, engaging preferred sets of ARCs in recruiting multivalent tankyrase binders such as AXIN ^48^. The TBM-binding pocket contains several binding hotspots (Figure 1B). An “arginine cradle” forms the binding site for the TBM’s essential arginine residue at position 1; the “central patch” provides an infrastructure for diverse interactions, including hydrophobic contacts with a small hydrophobic residue at TBM position 4 and contact sites for the residue at TBM position 5; and the “aromatic glycine sandwich”, where an essential glycine at TBM position 6 is sandwiched between two aromatic residues ^4^.

Mutation of the TBM binding sites in the ARCs abrogates tankyrase’s ability to drive Wnt/β-catenin signaling ^9^. As a further proof of concept for the feasibility of targeting tankyrase via the ARCs, a sequence-optimized ^4^, cell-permeating stapled TBM peptide can compete with AXIN for tankyrase binding and suppress the Wnt-induced expression of a β-catenin-responsive reporter gene in HEK293 cells ^50^. Given the uniqueness of ARCs within the PARP family and their strong degree of conservation across both TNKS and TNKS2, interfering with substrate binding would provide high target specificity and inhibition of both TNKS and TNKS2, many of whose functions are redundant ^6,51^.

Herein, we report the identification and characterization of fragments that bind to tankyrase ARCs at the same site as the TBM peptides. The identified fragments provide a starting point for the development of potent, cell-active tankyrase substrate binding antagonists.

## Results

### Essentiality of the TBM arginine residue

We first considered a peptidomimetic approach to develop TBM peptides into more potent, stable and drug-like competitors of the ARC:TBM interaction. Given the anticipated impairment of cell permeability by the N-terminal TBM arginine, we investigated whether the guanidine group could be substituted. To prioritise synthesis efforts, we followed an *in-silico* docking approach ^52^, exploring the importance of the positive charge and hydrogen bonding interactions, linker lengths/flexibility and side chain geometry (see Supplementary Materials and Methods for details). From commercially available side chain alternatives, we identified five potential candidates for R replacements: 1H-imidazole-5-pentanoic acid, 1H-imidazole-1-pentanoic acid, 7-aminoheptanoic acid, D-arginine and L-citrulline (Supplementary Figure 1). We next synthesised 3BP2 TBM octapeptides, incorporating the five arginine substituents at position 1, followed by fluorescence polarization (FP) assays to assess competition of the peptides with a Cy5-labelled TBM peptide probe (Supplementary Figure 1). We used a 16-mer TBM peptide (LPHLQ**RSPPDGQS**FRSW, W introduced to measure A_280_) derived from the model substrate 3BP2, a signaling adapter protein ^4^, as a positive control for a competitor, and a corresponding non-binding TBM peptide bearing a glycine-to-arginine substitution at position 6 ^4^ as a negative control. Whilst we observed no binding for the G6R negative control, we measured an IC_50_ of 22 μM for the 3BP2 16-mer peptide (Supplementary Figure 1). The 8-mer **RSPPDGQS** TBM peptide displayed an IC_50_ of 34 μM. Substituting L-arginine for D-arginine caused a five-fold drop in potency to 175 μM, highlighting the importance of side chain geometry. Both imidazole moiety peptides displayed IC_50_ values in the 500 μM range. The 7-aminoheptanoic acid and citrulline peptides showed poor competition and precipitation at high concentrations (Supplementary Figure 1). In conclusion, these observations demonstrated that the essential arginine residue of the TBM cannot be easily substituted.

### Primary fragment screens

Given the anticipated challenges associated with replacing the TBM arginine residue, we pursued a fragment screening strategy to sample a wide range of chemical space, with the aim of finding novel, ligand-efficient small molecules that target tankyrase ARCs and to identify alternative binding ‘hotspots’ away from the “arginine cradle” of the TBM binding site. We screened The Institute of Cancer Research (ICR) fragment library ^53^ in parallel against TNKS2 ARC5 using differential scanning fluorimetry (DSF), and TNKS2 ARC4 using ligand-observed nuclear magnetic resonance (NMR) spectroscopy techniques. We used the 3BP2 TBM peptide and its non-binding mutant variant as positive and negative controls, respectively.

### Primary fragment screening by DSF

Our pilot studies showed that among all TNKS2 single ARCs that we could produce recombinantly (ARCs 1, 4 and 5) ^49^, ARC5 displayed the lowest melting temperature (T_m_) and the largest shift in melting temperature upon addition of the 3BP2 TBM peptide (ΔT_m_) (Figure 2A). Therefore, we chose TNKS2 ARC5 for screening by DSF, anticipating the largest signal window for measuring changes in T_m_ upon fragment binding. DMSO concentrations up to 10% of total sample volume had a negligible effect on TNKS2 ARC5 T_m_ (Supplementary Figure 2A). We explored the stabilization of TNKS2 ARC5 by the abovementioned TBM peptide derivatives and found a good correlation between the DSF data and the FP data obtained with TNKS2 ARC4, further demonstrating the suitability of the DSF assay (Supplementary Figure 2B).

**Figure 2.**
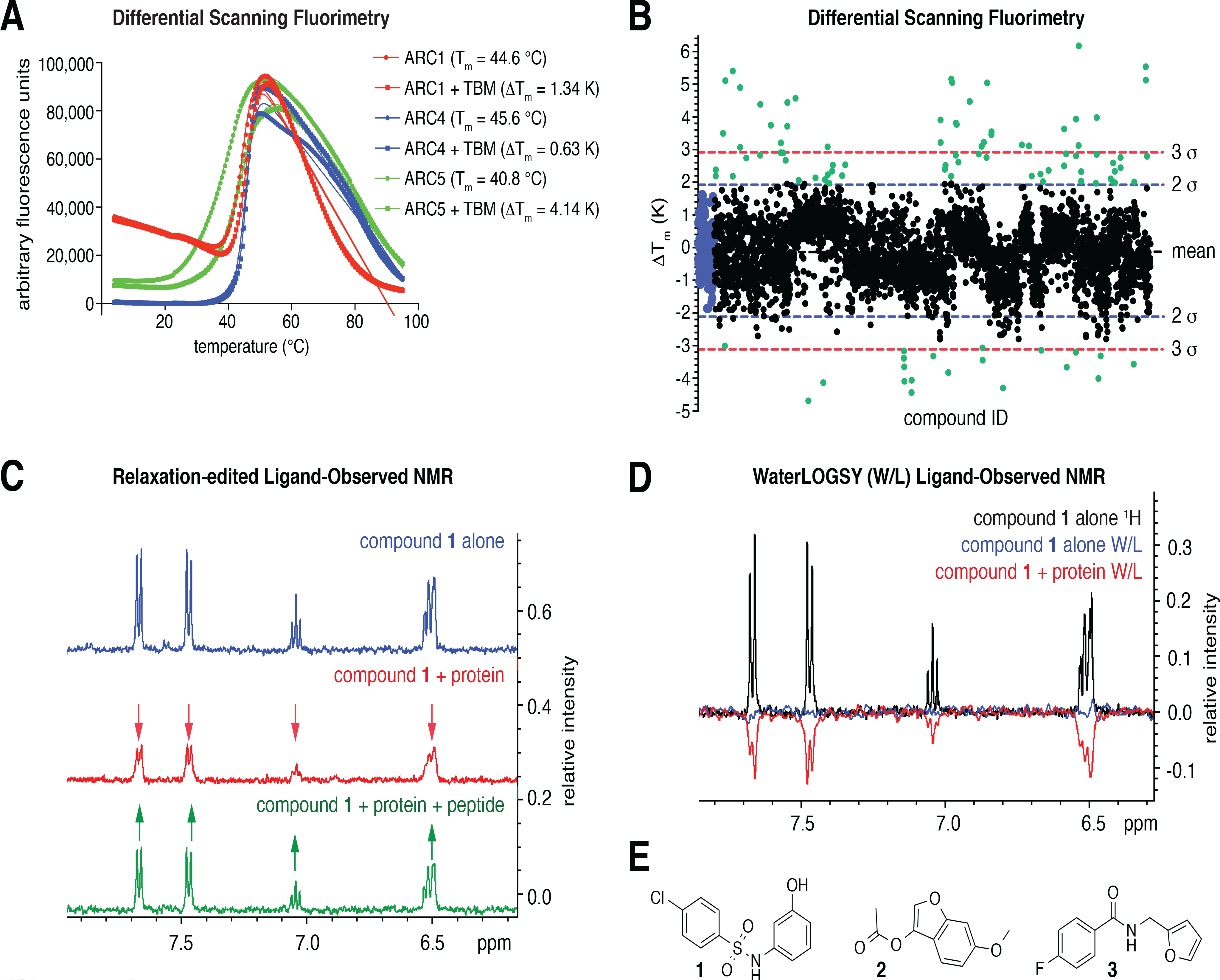
**(A)** Differential scanning fluorimetry (DSF, a.k.a. ThermoFluor) assessment of TNKS2 ARCs 1, 2 and 5 shows that TNKS2 ARC5 is the least stable among these ARCs and experiences the highest degree of thermal stabilization upon 3BP2 TBM peptide binding. **(B)** Fragment screen against TNKS2 ARC5 by DSF, showing compound ID vs. ΔT_m_ (from the IP method) for both replicates. DMSO-only controls are colored blue; hit fragments are colored green. Lines correspond to the mean, and 2 or 3 standard deviations outside the mean. **(C)** Example of relaxation-edited spectra for hit compound **1**. Signals are reduced in the presence of protein (red), indicating ARC binding, and recovered upon TBM peptide addition (green), indicating competition. **(D)** Example of waterLOGSY spectra for hit compound **1**, showing a negative NOE signal when protein is added (red). Buffer (HEPES) signals were phased as positive peaks in our waterLOGSY spectra. **(E)** Structures of hit compounds from the DSF screen that were competitive with a TBM peptide by relaxation-edited ligand-observed NMR.

We screened 1869 compounds in duplicate at a concentration of 500 μM, which we considered a reasonable compromise between having a sufficiently high concentration to identify weak binders while minimizing the likelihood of false positives through fragment precipitation and aggregation, and non-specific binding. The final DMSO concentration was 5%. We calculated melting temperatures using both the inflection point and the maximum peak of first derivative data methods. Both methods generally agreed, unless the melt curve was biphasic or misshapen, with slightly lower variability for the first derivative T_m_ determination method (see Materials and Methods for experimental and analysis details).

The melting temperature for the unbound ARC (T_m, 0_) was determined from the mean of 12 reference melting curves per plate, with 5% DMSO only. We calculated the change in melting temperature (ΔT_m_) by subtracting the mean T_m, 0_ from T_m, compound_. We tested compounds in duplicate, defining fragments as hits if they conferred a ΔT_m_ outside two standard deviations (2σ) from the mean, in one or both replicates. To check for consistency between plates, we ran triplicate peptide controls; however, we excluded peptide ΔT_m_ values from the calculation of the mean ΔT_m_ to avoid skewing the results. We observed mean ΔT_m_ values (IP/1^st^ derivative methods) of −0.127 / −0.331 K and σ as 0.997 / 1.19 K. A hit cut-off of 2σ gave absolute shifts of +1.87 / +2.05 K for compounds that stabilised and −2.13 / −2.71 K for compounds that destabilised the ARC (Figure 2B).

We assessed the robustness of the assay for screening. The standard deviation for both the DMSO-only (T_m, 0_) and 3BP2 peptide positive control (T_m, peptide_) melting temperatures across all plates was approximately 1 K, indicating that any shifts below 1 K may be attributable to noise. We calculated the Z factor (Z’) using the mean melting temperature and σ for the whole fragment library, with the DMSO-only samples as the baseline and 3BP2 TBM peptide samples as positive controls. A value of Z’ = 0.9 was obtained, indicating that the assay was robust.

We prioritised hits that stabilised TNKS2 ARC5 if they had a change in melting temperature of greater than 1.8 K in at least one replicate (both ΔT_m_(IP) and ΔT_m_(1st derivative) methods of analysis), and de-prioritised those that showed a significant discrepancy between ΔT_m_(IP) and ΔT_m_(1st derivative) values, indicating an unusual melting curve shape. Negative shift hits that destabilized the protein were only taken forward if they were significant in both replicates, and with both methods of ΔT_m_ analysis. (The higher stringency was applied as molecules that destabilize the protein can be harder to advance and develop into lead-like compounds ^54,55^.) We thus progressed 56 hits into validation assays. Of these, 48 conferred a positive thermal shift and stabilized TNKS2 ARC5; 8 had a negative thermal shift and destabilized the protein. We next assessed compound purity and structural integrity of hits from the DSF screen by liquid-chromatography-mass-spectrometry (LC-MS) and measured their solubility by NMR. Five compounds failed the LC-MS quality control, and eight were of insufficient solubility by NMR (<100 μM in aqueous buffer with 5% DMSO). Two compounds didn’t contain any aromatic protons (required for our NMR solubility assay), and an additional two were no longer commercially available for re-purchase. A total of 17 of compounds were therefore excluded from further analysis. An *in-silico* pan assay interference compounds (PAINS) screen was applied to the hit fragments to highlight any possible issues in carrying the hits forward ^56^. No compounds were flagged as problematic in the PAINS screen.

### Fragment binding validation for DSF hits

39 hits from the DSF screen were suitable for follow-up by ligand-observed NMR methods. We re-purchased fragments and performed T2 relaxation-edited (CPMG-edited) and waterLOGSY experiments for each fragment with TNKS2 ARC5. We explored saturation transfer difference (STD) NMR, also using TNKS2 ARC5, but the assay was not sensitive enough to produce a reliable binding signal, likely due to the relatively small size of a single ARC protein (data not shown). High ligand concentrations were required to achieve sufficient signal, which in turn could lead to false-positive hits due to non-specific binding.

Using the relaxation-edited assay, we tested each fragment in three independent measurements (Figure 2C), unless we obtained two negative results (non-binding) in the first two experiments. We next used waterLOGSY to further evaluate compounds that showed a substantial decrease (>15% reduction in peak integrals) upon protein addition in at least one out of three relaxation-edited experiments. We classified fragments with a negative NOE signal in waterLOGSY as binders (Figure 2D). As a negative NOE for the compound-only sample could indicate aggregation, we flagged these compounds as potentially problematic. We identified 14 fragments that bound to TNKS2 ARC5 by both relaxation-edited and waterLOGSY methods (0.78% hit rate). We next tested whether binding of these fragments occurred competitively with the 3BP2 TBM peptide, and also if they bound to TNKS2 ARC4, as competition with peptide binders at various different ARCs will be a prerequisite for an efficient substrate binding antagonist. Three fragments (**1**, **2**, **3**) bound to both TNKS2 ARC4 and ARC5 and were competitive with the TBM peptide (0.16% hit rate) (Figure 2E). Three further fragments also bound to both ARCs, but were not TBM competitive by NMR.

### Primary fragment screening by NMR

The hit rate for compounds confirmed to bind as evaluated by NMR was relatively low, at 0.78%, and only 0.16% for fragments competitive with a TBM peptide. Different screening assays often give distinct hit fragments ^57^. There is no consensus on the most appropriate assays to use for fragment screening, especially against challenging targets such as protein-protein interactions. Often several orthogonal methods are used in series to narrow down fragment hits, or a combination of biophysical and biochemical assays to exclude false positives and identify binders that modulate protein activity ^53^. We therefore carried out an additional primary screen using T2 relaxation-edited ligand-observed NMR on TNKS2 ARC4, using a subset of the ICR fragment library that was compatible with NMR. We screened 1100 compounds in pools of four structurally dissimilar molecules with non-overlapping proton resonances (Figure 3A, B) ^58^. We split the top 100 hits into two groups for individual re-testing: those with a signal change ≥39% (3σ, 35 compounds), and those with a signal change of 26 – 39% reduction (2 – 3σ, 65 compounds) (Figure 3A). We tested fragments of the first hit group (>3σ) individually using the T2 relaxation-edited NMR assay. We tested those of the second hit group (2 – 3σ) in a waterLOGSY NMR experiment, reasoning that this may rescue any genuine binders with a relatively small signal in the relaxation-edited assay. Nine out of 35 compounds from hit set 1 displayed a significant intensity change (≥26% reduction) upon protein addition when tested individually. We confirmed seven out of 65 compounds from hit set 2 to bind in the waterLOGSY assay. We tested these 16 compounds in further T2 relaxation-edited and waterLOGSY experiments, and in competition with the 3BP2 TBM peptide. Of the 16 compounds, two (**4** and **5**) were competitive with the TBM peptide by relaxation-edited NMR; one compound (**5**) also showed peptide competition by waterLOGSY (Figure 3C).

**Figure 3.**
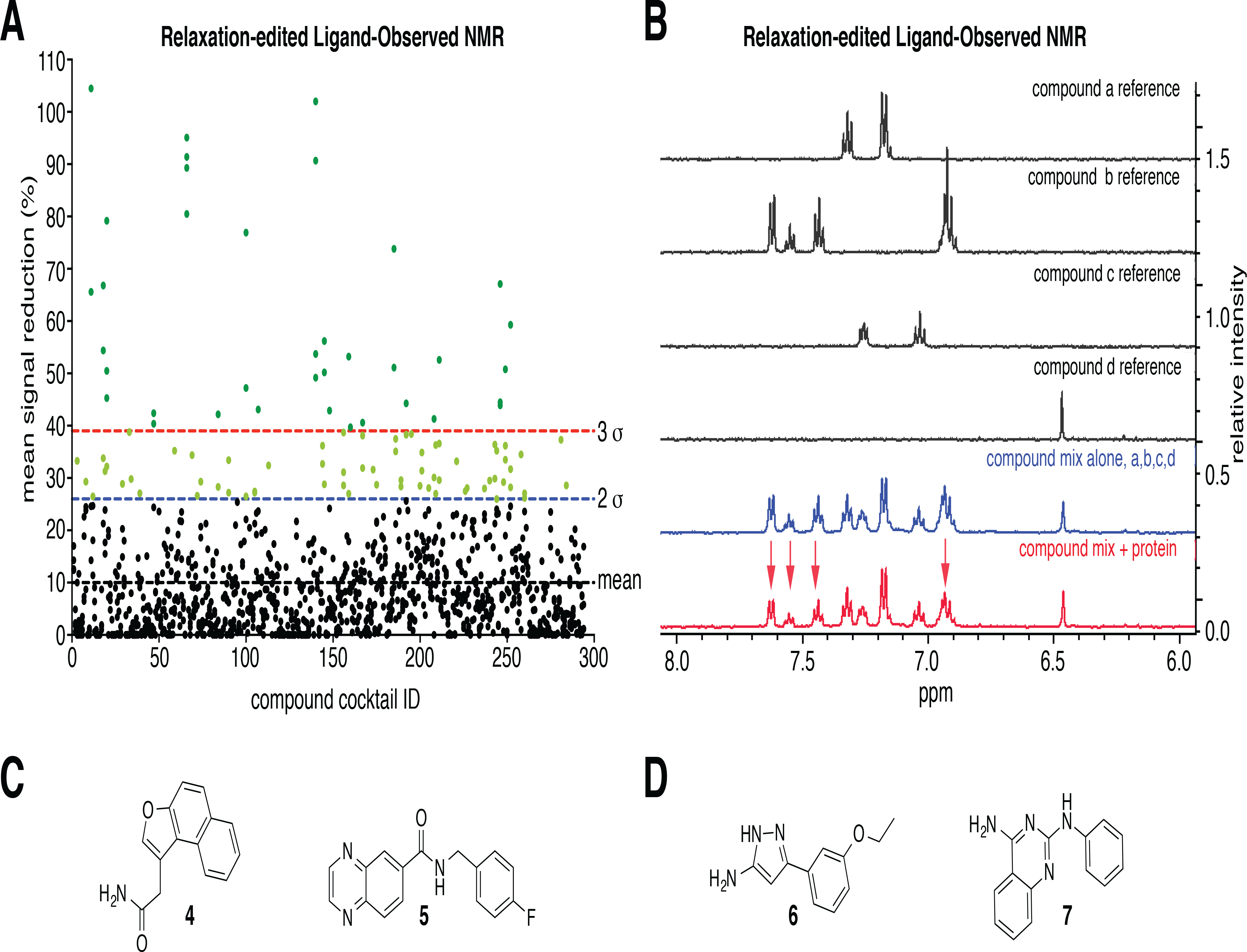
**(A)** Summary of ligand-observed NMR screen, showing percentage of signal change vs. compound cocktail ID. Lines correspond to the mean, and 2 or 3 standard deviations outside the mean. **(B)** Example data for a cocktail of 4 compounds, containing one hit (compound b) and three non-binding fragments. **(C)** Structures of hit fragments uniquely identified in the NMR screen, which were TBM-competitive by relaxation-edited NMR. **(D)** Structures of compounds that were identified as hits in both the DSF and NMR primary screens.

In addition to the hits identified uniquely by either the DSF or NMR, the two orthogonal screens also shared two common hits (compounds **6** and **7**, Figure 3D). Retrospective analysis of the DSF screening data revealed that compound **5** was excluded during the DSF analysis due to poorly shaped, biphasic melt curves. This resulted in large discrepancies between ΔT_m_ values calculated by the inflection point (1.6 K) and 1^st^ derivative methods (8.1 K) when the second peak was used to calculate T_m_. We also discounted other compounds for poor melt curve shape; however, none of these were identified in the orthogonal NMR-based screen.

### Fragment hit validation and K_d_ determination

We next tested validated hits from both the DSF and NMR screens (17 compounds in total) against TNKS2 ARC4 using protein-observed NMR. We used the 3BP2 TBM peptide as a positive control. To directly identify fragment binding sites on the ARC, we performed a full backbone and partial side-chain assignment of TNKS2 ARC4, doubly labelled with ^15^N and ^13^C. The assignment details will be reported elsewhere (Zaleska *et al.*, 2019) ^69^. The spectra showed significant chemical shift perturbations (CSPs) induced by the peptide, indicative of peptide binding in both a fast and slow kinetic regime (Supplementary Figure 3). Interestingly, among the residues that constitute the TBM binding site on the ARC, residues that exhibited the slow-exchange binding mode are part of the “central patch” (D521, S527, F532, D556, L560, H564, N565, S568) and the “aromatic glycine sandwich” (G535, Y536, Y569); a single residue from the “arginine cradle” (F593) displayed slow exchange. This suggests that these areas form key TBM:ARC interaction hotspots. Residues that exhibited the fast-exchange binding mode were from the “arginine cradle” (D589, W591, E598) and the “C-terminal contacts” (H571, K604). The different binding regimes may distinguish primary interaction hotspots that are engaged robustly when a peptide is first recruited (slow exchange) from secondary binding sites in the ARC that become occupied once primary binding hotspots are engaged; these may be more dynamic (fast exchange).

We titrated compounds against ^15^N-labeled TNKS2 ARC4, initially at protein:compound ratios of 1:1 and 1:3. Two fragments (**3** and **5**) induced significant CSPs (data not shown); we used these to perform an eight-point titration and observed concentration-dependent CSPs (Supplementary Figure 4A, B). We confirmed that the CSPs were not caused by pH changes during the titration by measuring the pH of 3 mM peptide and fragment stocks in assay buffer. Consistent with TBM-competitive binding of fragments **3** and **5**, we identified several peaks that shifted in both fragment and 3BP2 TBM peptide titrations. Compound **5** caused more significant perturbations than compound **3**, and they all exhibited a fast-exchange regime (Supplementary Figure 4C). Peaks that moved significantly (CSPs > Δδ_tot_ + 2σ) upon addition of compound **5** are part of the “central patch” and “aromatic glycine sandwich” (S527, F532, G535, Y536, N565; see Supplementary Figure 4C). However, the solubility of fragments **3** and **5** in assay buffer limited the maximum concentration achievable, and so complete saturation was not reached, which confounded affinity measurements and more extensive analyses of the fragment binding sites by protein-observed NMR.

### Fragment analog SAR

We next sought close structural analogs of hit fragments **3** and **5** to gain early insights into structure-activity relationships (SAR) and binding modes of hit fragments. We initially tested the analogs using both relaxation-edited and waterLOGSY NMR assays against TNKS2 ARC4 (Table 1), followed by protein-observed NMR if binding was observed in both ligand-observed NMR experiments. Compounds displaying negative NOE signals in the waterLOGSY assay upon protein addition were classed as binders; however, compounds that displayed negative NOE signals in the absence of protein were flagged as potential aggregators. Compounds that displayed no NOE signal were classed as non-binders.

**Table 1.**
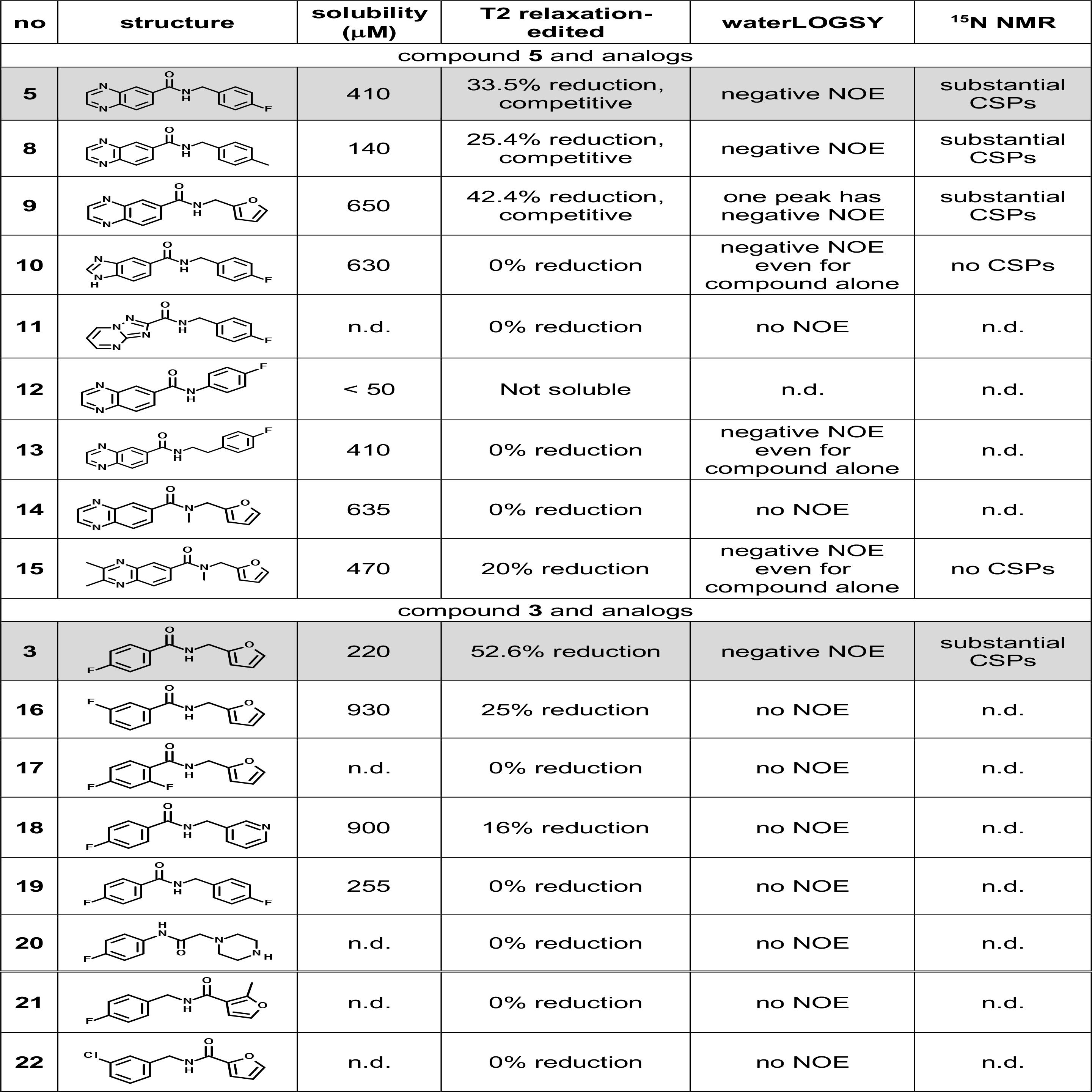
Analogs of compounds **3** and **5** tested by ligand-observed and protein-observed NMR. Compounds **17, 18 and 20** – **22** were already present in the fragment library and were therefore not re-tested. Rows shaded grey are the original screening hits. “n.d.”, not determined.

| no | structure | solubility<br>(μM) | T2 relaxation-edited | waterLOGSY | $^{15}\text{N}$ NMR |
| --- | --- | --- | --- | --- | --- |
| compound **5** and analogs |  |  |  |  |  |
| **5** |  | 410 | 33.5% reduction,<br>competitive | negative NOE | substantial<br>CSPs |
| **8** |  | 140 | 25.4% reduction,<br>competitive | negative NOE | substantial<br>CSPs |
| **9** |  | 650 | 42.4% reduction,<br>competitive | one peak has<br>negative NOE | substantial<br>CSPs |
| **10** |  | 630 | 0% reduction | negative NOE<br>even for<br>compound alone | no CSPs |
| **11** |  | n.d. | 0% reduction | no NOE | n.d. |
| **12** |  | < 50 | Not soluble | n.d. | n.d. |
| **13** |  | 410 | 0% reduction | negative NOE<br>even for<br>compound alone | n.d. |
| **14** |  | 635 | 0% reduction | no NOE | n.d. |
| **15** |  | 470 | 20% reduction | negative NOE<br>even for<br>compound alone | no CSPs |
| compound **3** and analogs |  |  |  |  |  |
| **3** |  | 220 | 52.6% reduction | negative NOE | substantial<br>CSPs |
| **16** |  | 930 | 25% reduction | no NOE | n.d. |
| **17** |  | n.d. | 0% reduction | no NOE | n.d. |
| **18** |  | 900 | 16% reduction | no NOE | n.d. |
| **19** |  | 255 | 0% reduction | no NOE | n.d. |
| **20** |  | n.d. | 0% reduction | no NOE | n.d. |
| **21** |  | n.d. | 0% reduction | no NOE | n.d. |
| **22** |  | n.d. | 0% reduction | no NOE | n.d. |

Substituting the para-fluorine of compound **5** for a methyl group retained binding (**8**), as did replacement of the entire Ar-F group with a furan (**9**). Analog **9** showed increased solubility over original hits **3** and **5**. Contracting the quinoxaline ring by one carbon atom to a benzimidazole (**10**) abrogated binding. Substitution of the quinoxaline moiety by a triazolopyrimidine (**11**) also abrogated binding. Shortening the amide linker by one carbon (**12**) limited solubility. Increasing the linker length by one carbon (**13**) abolished binding in the relaxation-edited NMR assay, however, gave a strong waterLOGSY signal. We also observed a strong waterLOGSY signal for compound **13** in the absence of protein, indicating that this compound may aggregate. Methylating the amide nitrogen of **9** was not tolerated (**14**). Additionally substituting the quinoxaline ring at positions 2 and 3 with methyl groups **(15)** showed binding by relaxation-edited ligand-observed NMR; however, the waterLOGSY data suggested compound aggregation, and no CSPs were observed in protein-observed NMR. In summary, we demonstrated TNKS2 ARC4 binding activity of several quinoxaline analogues of compound **5**, confirming this hit series and showing that the Ar-F group of **5** could be readily substituted.

For the benzamide fragment (**3**), moving the fluorine atom from the para to the meta position **(16)** or adding an ortho-fluorine (**17**) abolished binding. Substitution of the furan for a pyridine (**18**) or Ar-F moiety (**19**) was not tolerated. Reversing or rearranging the amide linker (**20, 21**, **22**) whilst simultaneously changing the furan for a piperidine (**20**), adding a methyl group to the furan ring (**21**) or substituting the para-fluorine for a meta-chlorine (**22**) were also not tolerated.

Compound **9** (Table 1, Figure 4A) combined features of both fragments **3** and **5**, namely the quinoxaline group of fragment **5**, the amide linker shared by both fragments and the furan group of fragment **3**. We observed that in the relaxation-edited NMR experiments, peaks corresponding to the quinoxaline displayed a larger reduction upon ARC addition than peaks attributed to furan (Figure 4A). This suggested that the quinoxaline moiety more substantially contributes to the binding, and several analogues of the quinoxaline hit were confirmed to bind to TNKS2 ARC4.

**Figure 4.**
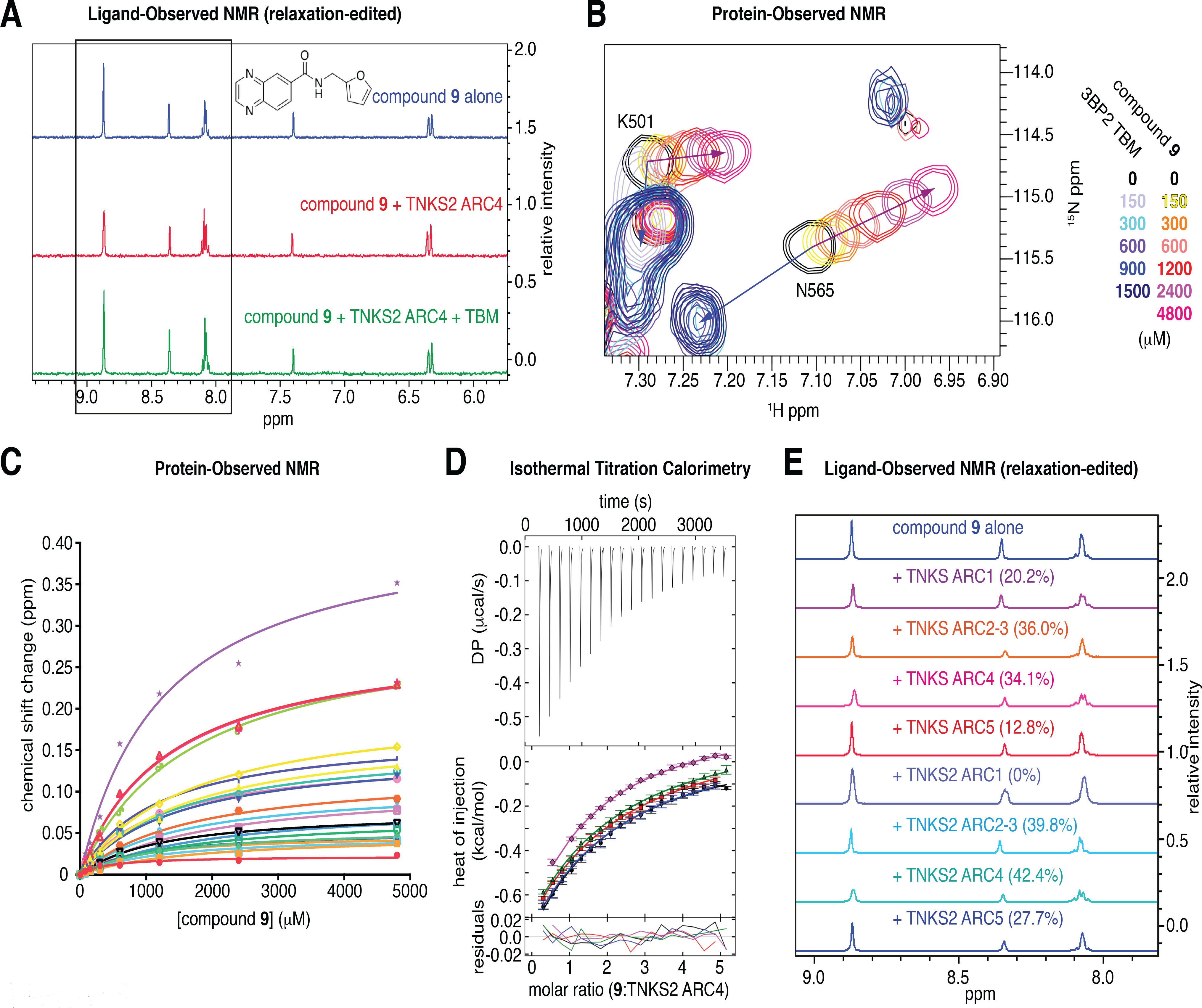
**(A)** Relaxation-edited NMR for compound **9**, showing the largest reduction in peak height for the quinoxaline protons (boxed), indicating that the majority of binding can be attributed to the quinoxaline moiety. **(B)** ^1^H-^15^N HSQC NMR spectrum, showing the chemical shift perturbation upon TBM peptide or compound **9** addition. **(C)** K_d_ estimate of compound **9** by plotting the chemical shift perturbation of peaks that moved in a concentration-dependent manner. **(D)** ITC for the titration of compound **9** (5 mM) into TNKS2 ARC4 (200 µM). The K_d_ for compound **9** was calculated to be 1200 ± 380 μM (global analysis of n = 5 independent experiments). **(E)** Compound **9** binding to TNKS and TNKS2 ARCs was assessed by relaxation-edited NMR. Total reductions in peak area upon ARC addition are indicated.

### Fragment binding affinity

The increased solubility of compound **9** compared to compound **5** allowed complete saturation in a protein-observed NMR titration experiment against TNKS2 ARC4, yielding an apparent K_d_ of 1050 μM (Figure 4B, C). We next used isothermal titration calorimetry (ITC) to confirm the compound **9**:TNKS2 ARC4 binding affinity, titrating the fragment (5 mM, 1% DMSO) into TNKS2 ARC4 (200 μM, 1% DMSO). A global analysis of 5 experiments confirmed the affinity to be in the region of 1 mM (1200 ± 380 μM) (Figure 4D, Table 2).

**Table 2.**
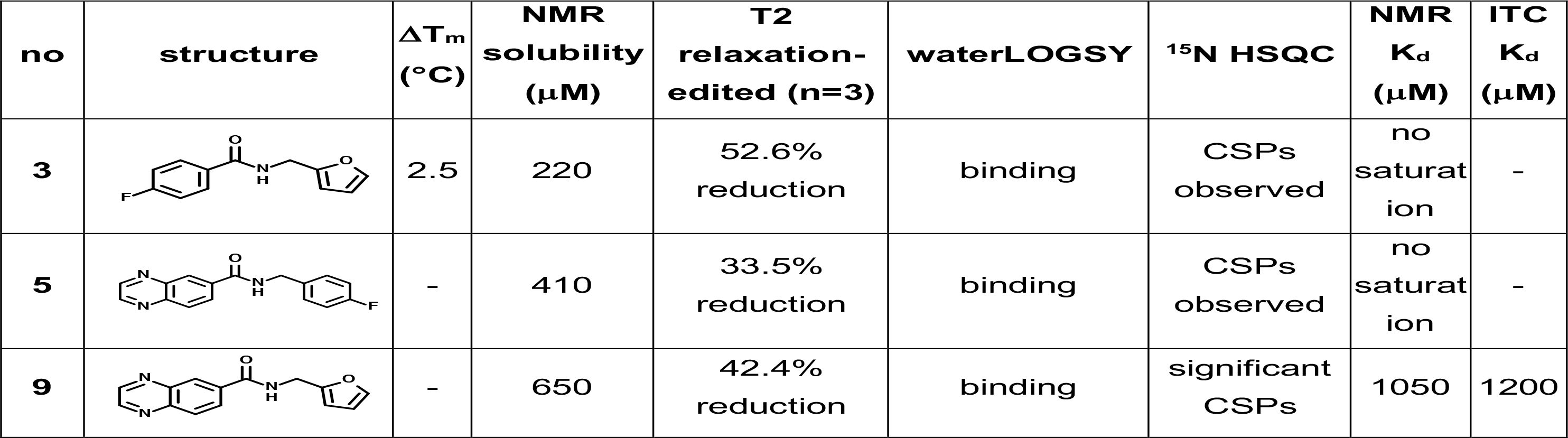
Summary of data for fragments confirmed to bind to TNKS2 ARC4 by protein-observed NMR.

| no | structure | $\Delta T_m$<br>(°C) | NMR<br>solubility<br>(μM) | T2<br>relaxation-edited (n=3) | waterLOGSY | $^{15}\text{N}$ HSQC | NMR<br>$K_d$<br>(μM) | ITC<br>$K_d$<br>(μM) |
| --- | --- | --- | --- | --- | --- | --- | --- | --- |
| **3** |  | 2.5 | 220 | 52.6%<br>reduction | binding | CSPs<br>observed | no<br>saturat<br>ion | - |
| **5** |  | - | 410 | 33.5%<br>reduction | binding | CSPs<br>observed | no<br>saturat<br>ion | - |
| **9** |  | - | 650 | 42.4%<br>reduction | binding | significant<br>CSPs | 1050 | 1200 |

### Fragments bind to multiple TNKS and TNKS2 ARCs

The anticipated functional redundancy between ARCs and the existence of two tankyrase paralogs will require efficient substrate binding antagonists to ideally bind all TBM-binding ARCs of both TNKS and TNKS2. The high conservation of the peptide-binding pocket suggests that this goal should be achievable ^4^. We tested binding of compound **9** to all TNKS and TNKS2 ARCs using the relaxation-edited NMR assay (Figure 4E, Table 3). Compound **9** bound all ARCs, with the exception of TNKS2 ARC1. Given the invariant residue infrastructure of the peptide-binding pocket in ARC1 of TNKS and TNKS2, this observation is difficult to reconcile with available structural information. It is possible that the presence of glycine-sandwiching phenylalanine rather than tyrosine residues in ARC1 of both tankyrases, paired with the presence of a phenylalanine in TNKS2 (F29^TNKS2^) as opposed to a leucine in TNKS (L187^TNKS^) confers this differential behaviour. F29^TNKS2^/L187^TNKS^ sit in an extended hydrophobic pocket adjacent to the glycine sandwich, which may participate in fragment binding. Whilst the low affinity of the current fragments may sensitize them to small differences between the TBM-binding ARCs, further developed molecules will need to be engineered to be resistant to such variability.

**Table 3.** Pan-ARC binding activity of compound **9**, tested by ligand-observed NMR.

| ARC construct | T2 relaxation-edited | waterLOGSY |
| --- | --- | --- |
| TNKS ARC1 (178 – 336) | 20.2% reduction | negative NOE signal |
| TNKS ARC2-3 (331 – 645) | 36.0% reduction | negative NOE signal |
| TNKS ARC4 (646-807) | 34.1% reduction | negative NOE signal |
| TNKS ARC5 (799-958) | 12.8% reduction | negative NOE signal |
| TNKS2 ARC1 (20-178) | 0% | no NOE |
| TNKS2 ARC2-3 (173 – 487) | 39.8% reduction | negative NOE signal |
| TNKS2 ARC4 (488-649) | 42.4% reduction | negative NOE signal |
| TNKS2 ARC5 (641-800) | 27.7% reduction | negative NOE signal |

### *In-silico* modeling of fragment hotspots

We next sought to determine the fragment binding site on the ARC. To gain insights into plausible fragment binding sites and identify potential hotspots, we undertook an *in-silico* fragment binding experiment by computational solvent mapping using the FTMap program ^59,60^. FTMap identifies pockets where several different small organic molecule probes bind and cluster together; these consensus sites represent potential hotspots for fragment binding. We docked a set of 16 probe molecules into the crystal structure TNKS2 ARC4, from the ARC4:3BP2 TBM co-crystal structure ^4^. FTMap identified ten areas of probe clustering, eight of which overlapped with the known peptide-binding groove (Figure 5A). The lowest-energy consensus site, and hence most ligandable pocket identified, was the primarily hydrophobic “central patch” adjacent to the “glycine sandwich”. The second most ligandable site predicted was the “arginine cradle”. These predicted hotspots coincide with the experimentally determined hotspots for TBM peptide binding, based on structural data, site-directed mutagenesis and an amino acid scan of the 3BP2 TBM ^4,61^. One hotspot was detected in an extension to the “central patch”, suggesting that it may be possible to grow fragments in a way that utilises this extended pocket. Of note, this “central patch extension” is occupied by a glycerol molecule in the TNKS2 apo-ARC4 crystal structure ^4^. The potential fragment binding sites located by FTMap coincide with pockets identified using the program Pocasa, which performs a geometric search based on a three-dimensional grid and rolling probe sphere ^62^ (Figure 5B). The “central patch” and “central patch extension” were the highest-ranked pockets, followed by the “arginine cradle”, with volumes/volume depth values of 126/289, 46/108 and 26/73, respectively.

**Figure 5.**
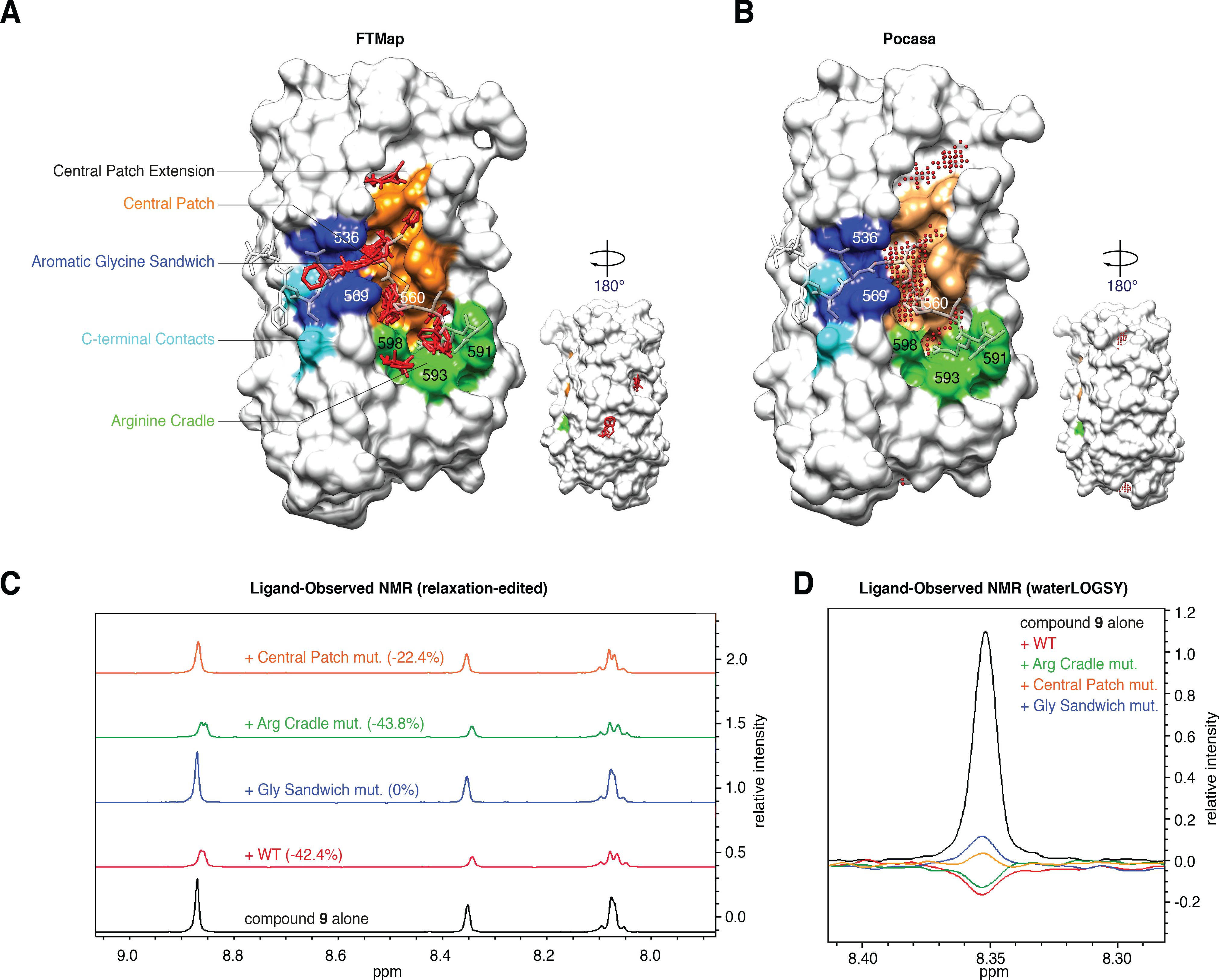
**(A)** Eight fragment binding hotspots on TNKS2 ARC4 predicted by FTMap are in the TBM peptide-binding site on the ARC or in close vicinity. FTMap analysis was done on TNKS2 ARC4 from the ARC4:3BP2 co-crystal structure ^4^ (3TWR). Key residues of the peptide binding site are color-coded as in Figure 1B. The TBM peptide from 3BP2 is overlaid in transparent stick representation. The minimum energy hotspot found is in the “central patch” adjacent to the “glycine sandwich”. **(B)** Pocket identification on TNKS2 ARC4 (from the ARC4:3BP2 co-crystal structure, 3TWR) using the Roll algorithm implemented in Pocasa ^62^. The three top-ranking pockets are part of the “central patch”, the “central patch extension” and the “arginine cradle”. **(C)** Relaxation-edited NMR of compound **9** with TNKS2 ARC4 peptide binding site mutants ^4^. Mutation of the “aromatic glycine sandwich” abolishes binding of the compound, while binding is unaffected by mutation of the “arginine cradle”. Mutated residues, numbered in (A) and (B), were as follows: “arginine cradle”, WFA591/593/598AAA; “central patch”, L560W; “aromatic glycine sandwich”, YY536/569AA. **(D)** WaterLOGSY NMR confirms that “glycine sandwich” and “central patch” mutations impair binding of compound **9**.

### Ligand-observed NMR with mutant TNKS2 ARC4 proteins

We used three previously designed TNKS2 ARC4 mutant variants ^4^ to explore potential binding determinants for fragment **9**. A WFE591/593/598AAA triple-mutation abrogates three key residues in the “arginine cradle”; YY536/569AA truncates two tyrosine residues that form the “glycine sandwich”, and L560W introduces a bulky residue into the “central patch” that sterically clashes with the TBM peptide (see Figures 5A, B for location of the mutated residues). We tested binding of compound **9** to the wild-type and mutant ARCs by ligand-observed NMR, using the relaxation-edited assay (Figure 5C, Table 4). While mutation of the “arginine cradle” had no effect on fragment binding, mutation of the aromatic residues sandwiching the TBM glycine (YY536/569AA) fully abrogated binding in the relaxation-edited NMR assay. Binding was impaired but not abolished for the “central patch” mutant variant (L560W). We confirmed these results in the orthogonal waterLOGSY assay (Figure 5D, Table 4).

**Table 4.** Ligand-observed NMR analysis to assess compound **9** binding to wild-type and mutant variants of TNKS2 ARC4.

| TNKS2 ARC4 constructs | T2 relaxation-edited | waterLOGSY |
| --- | --- | --- |
| compound alone | - | no NOE |
| wild-type | 42.4% reduction | negative NOE |
| arginine cradle<br>(WFE591/593/598AAA) | 43.8% reduction | negative NOE |
| central patch<br>(L560W) | 22.4% reduction | positive NOE |
| aromatic glycine sandwich<br>(YY536/569AA) | 0% | positive NOE |

### Fragment binding site mapping by protein-observed NMR

To directly identify the compound **9** binding site on the ARC, we analysed the titrations of compound **9** with ^15^N-labeled TNKS2 ARC4. The higher solubility of fragment **9**, compared to that of compounds **3** and **5**, meant that much larger CSPs could be achieved (Supplementary Figure 4C). At an ARC4:compound ratio of 1:16, close to signal saturation (see Figure 4C), we observed substantial CSPs for the following main-chain resonances: with CSPs above 2 σ from the mean CSP for S527, T528, F532, Y536, N565 and A566, and with CSPs within 1-2σ for A499, K501, D521, I522, L530, A534, G535 and L563 (Figure 6). All CSPs occurred in the fast-exchange regime and included those observed for compounds **3** and **5** (Figure 6, Supplementary Figure 4C). Mapping the CSPs onto the crystal structure of TNKS2 ARC4 bound to the TBM peptide from 3BP2 ^4^ revealed the substantial overlap of the fragment and TBM peptide binding site in the “aromatic glycine sandwich”, the “central patch” and residues in the close vicinity to these areas, in agreement with the mutagenesis studies (Figure 6A). In conclusion, compound **9** occupies the most ligandable pocket on the ARC and a major binding hotspot of TBM peptides.

**Figure 6.**
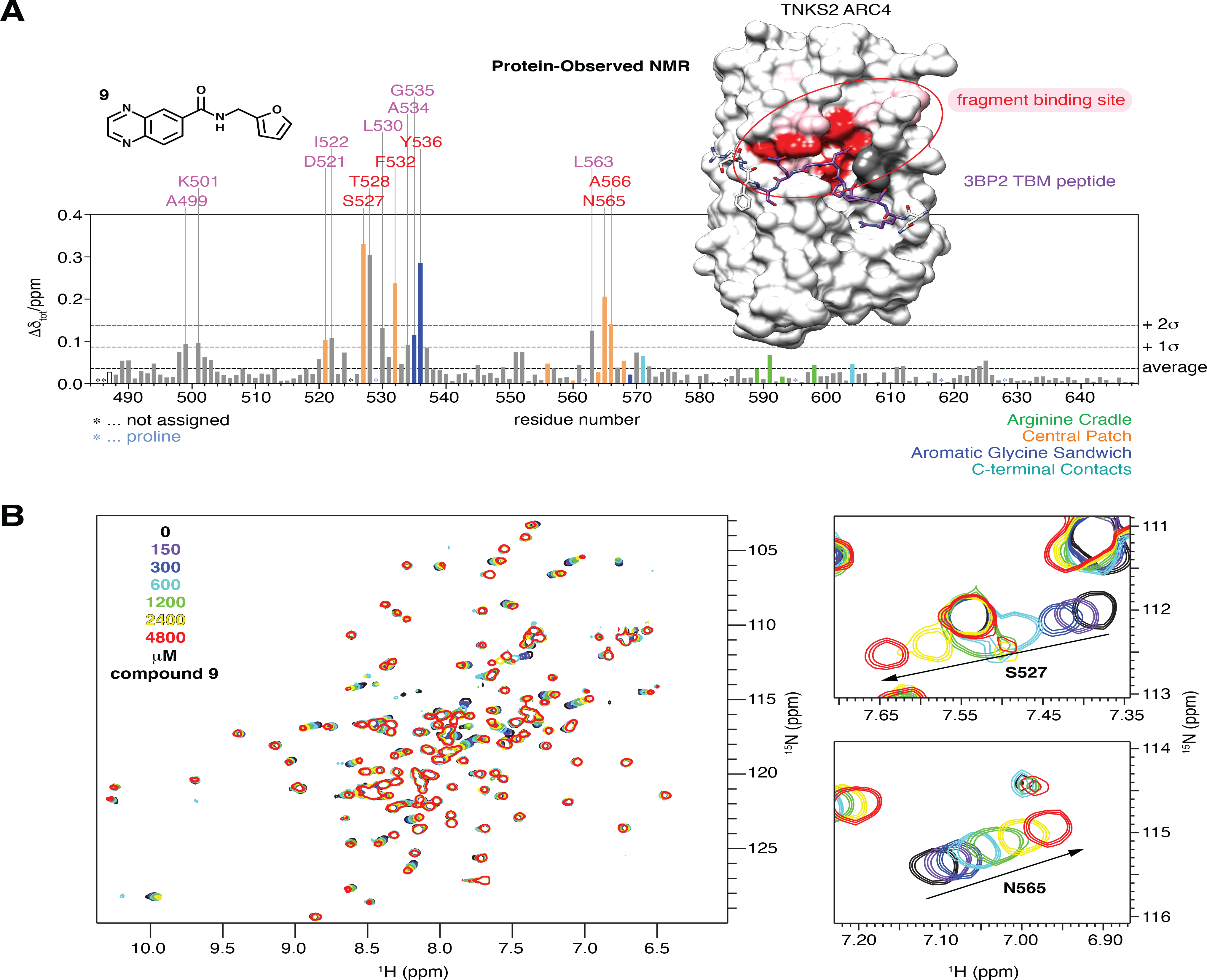
**(A)** Plot of CSPs in TNKS2 ARC4 (300 μM) induced by the addition of 16-fold excess (4800 μM) of compound **9** (see Figures 4B and C). Bars corresponding to residues known to bind the TBM ^4^ are color-coded as in Figures 1C, 5A and 5B. CSPs are mapped onto the surface representation of TNKS2 ARC4 bound to the TBM peptide of 3BP2 (shown in stick representation): the strongest perturbations (>2σ of average) are shown in red, weaker ones (>1σ and ≤2σ) in pink. The overlap of the TBM binding pocket and compound **9**-induced CSPs is clearly apparent. Unassigned residues are shown in dark gray. Prolines are shown in light blue (none on the peptide-binding face of the ARC). **(B)** Whole ^1^H-^15^N HSQC NMR spectrum of TNKS2 ARC4 and selected areas, showing the CSPs upon compound **9** titration.

## Conclusions and Discussion

Here we identify a quinoxaline-based set of fragments that bind to the substrate/protein-binding ARCs of tankyrase at the same site as TBM peptides. We show that the fragments bind in the “aromatic glycine sandwich” and “central patch” regions, major known hotspots of TBM binding. These fragments, even at their current affinities in the millimolar range, provide a potential starting point for the development of tool compounds to investigate the scaffolding roles of tankyrase, with the aim to validate whether tankyrase substrate binding antagonists are a viable approach to inhibiting tankyrase function.

Synthesising a set of TBM peptide variants as part of an initial peptidomimetic approach, we find that the essential, invariant arginine residue at TBM position 1 is challenging to replace with other groups that are less likely to impair cell permeability. Given that neither of the arginine substituents analysed here sufficiently preserved TBM binding, we took a fragment screening approach. Preceding *in-silico* analyses point to the “central patch” region as the top-ranking, potentially ligandable pocket of the ARC. Indeed, the identified fragment binding site overlaps substantially with the “central patch” and with the adjacent “aromatic glycine sandwich”, an anchor point for another invariant TBM residue, a glycine residue at TBM position 6. In protein-observed NMR studies, both the “central patch” and “aromatic glycine sandwich” coincide with TBM-peptide-induced CSPs in the slow-exchange kinetic regime. This further confirms their critical role in TBM binding and illustrates that the fragments indeed target key determinants of the ARC:TBM interaction.

Effective substrate binding antagonists will likely need to target all TBM-binding ARCs in both tankyrase paralogs. Given the high degree of conservation between TNKS and TNKS2 ARCs, and the nearly identical TBM-binding infrastructure between different ARCs ^4,48^, this appears feasible. We indeed observe multi-ARC-binding activity of compound **9**. The non-detectable binding of this fragment to TNKS2 ARC1 may be a consequence of its low affinity for tankyrase in general. Multi-ARC binding will need to be monitored as more potent compounds are developed.

Future studies will focus on the structure-based design of TBM competitors with increased affinity. Fully developed tankyrase substrate binding antagonists will enable the complex mechanisms of tankyrase to be probed in a wide range of its biological functions. In the longer term, substrate binding antagonists may be of potential therapeutic value as they offer an opportunity to block both catalytic and non-catalytic functions and may display pharmacodynamics that substantially differ from compounds targeting the tankyrase PARP catalytic domain.

## Materials and Methods

### Fragment screening using a thermal shift assay

For the screen, a C1000 thermal cycler (Bio-Rad) was used to record melt curves. SYPRO Orange was purchased as a 5000 × stock in DMSO from Sigma Aldrich. The ICR fragment library was available as 100 mM stocks in DMSO, and dispensed (25 nL) using an ECHO acoustic liquid handling system into white 384-well PCR plates (Framestar, 4titude). Wells were backfilled with DMSO (225 nL). Buffer (25 mM HEPES-NaOH pH 7.5, 100 mM NaCl, 2 mM TCEP, 2.75 μL) was added, followed by TNKS2 ARC5 (1 μL, 100 μM stock) and then SYPRO orange dye (1 μL, 25×). The plate was centrifuged after the addition of each reagent (1 min, 1000 × g). Final assay concentrations were as follows: TNKS2 ARC5 (20 μM); fragment (500 μM); SYPRO Orange dye (5×); DMSO (5% v/v) in a total volume of 5 μL. Peptide control wells (3BP2 TBM 16-mer, 200 μM, sequence LPHLQ**RSPPDGQS**FRSW with C-terminal tryptophan added for photometric concentration measurements, the N-terminus acetylated and the C-terminus amide-capped) were plated in triplicate, and there were 12 blank wells per plate, with DMSO only (250 nL, 5% v/v). Melt curves were recorded from 20-95 °C, with the temperature ramped by 0.5 °C every 15 s. Data were analysed using Vortex (Dotmatics) software, and the melting temperature was calculated from both the inflection point and the maximum peak of 1^st^-derivative data. Data points were excluded if the melt curve was poor, i.e. there was no fluorescence signal above baseline, there was high fluorescence intensity throughout, or the melt curve was shallow (<1000 rfu difference between baseline and peak maximum). The unbound melting temperature was determined from the mean of 12 reference melting curves, with 5% DMSO only. The change in melting temperature (ΔT_m_) was calculated by subtracting the mean T_m, 0_ from T_m, compound_. Compounds were tested in duplicate, and hits were defined as fragments that gave a ΔT_m_ outwith 2 σ from the mean in one or both replicates.

For the experiment shown in Figure 2A, ARC and 3BP2 TBM peptide concentrations were 20 and 200 μM, respectively, in 50 mM HEPES-NaOH pH 7.5, 100 mM NaCl, 2 mM TCEP and a total volume of 25 μL. Sypro Orange was added at 5×. Data were recorded from 4-95 °C, with the temperature ramped by 0.5 °C every 15 s, using a CFX384 thermal cycler (Bio-Rad). Data were analysed by non-linear regression in GraphPad Prism using a Bolzmann sigmoid with linear baselines. ΔT_m_ values were determined using the inflection point method.

### NMR experiments

A Bruker 500 MHz instrument, fitted with a 1.7 mm TXI microprobe was used for all ligand-observed and protein-observed NMR experiments, with 1.7 mm SampleJET NMR tubes (Bruker).

### Fragment solubility assay

Fragments were dispensed into a 384-well plate (250 nL of 100 mM stock in DMSO, 500 μM final concentration) using an ECHO acoustic dispenser. DMSO (2.25 μL) was added, then NMR buffer (47.5 μL, 25 mM HEPES-NaOH pH 7.5, 100 mM NaCl, 1 mM TCEP, 10% D_2_O). The plate was centrifuged (1 min, 1000 × g) before the solutions were transferred to 1.7 mm NMR tubes using a Gilson liquid handling system. ^1^H NMR spectra were recorded with the DMSO and water signals dampened. 100 μM caffeine was used as an external standard to quantify the ligand signals.

### T2 relaxation-edited NMR assay

Fragments were dispensed in duplicate into a 384-well plate (250 nL, 100 mM stock in DMSO). Wells were backfilled with DMSO (2.25 μL). Tankyrase ARC protein (47.5 μL, 20 µM in 25 mM HEPES, pH 7.5, 100 mM NaCl, 2 mM TCEP, 10% D_2_O) was added to the ‘protein’ samples; buffer only (47.5 μL) was added to the ‘compound-only’ samples. Solutions were transferred to 1.7 mm NMR tubes using a Gilson liquid handler. ^1^H NMR spectra were recorded, with the DMSO and water signals dampened. A relaxation spin filter was applied at 400 ms ^64^. Data were processed using Bruker Topspin 3.14. Lines were broadened with LB = 3.0, and the baseline was corrected between 6.0 – 10.0 ppm. The average integral for all peaks between 6.0 – 10.0 ppm was calculated, and the difference between compound-only and compound-plus-protein samples was compared. A reduction in signal integrals of ≥15% was classified as a hit. For competitive experiments, 3BP2 (100 μM) was added, and the spectra recorded and processed as above. The variability in signal reduction in the relaxation-edited experiment was previously determined as approximately ±10% (Liu *et al*., unpublished observations), so replicates were run to account for this variability, to ensure that compounds that had a weak reduction in signal that did not meet the arbitrary cut off were not erroneously excluded.

### WaterLOGSY NMR assay

Fragments were dispensed in duplicate into a 384-well plate (250 nL, 100 mM stock in DMSO). Wells were backfilled with DMSO (2.25 μL). Tankyrase ARC protein (47.5 μL, 20 µM in 25mM HEPES-NaOH pH7.5, 100 mM NaCl, 2 mM TCEP, 10% D_2_O) was added to the samples containing protein; buffer only (47.5 μL) was added to the compound-only samples. Solutions were transferred to 1.7 mm NMR tubes using a Gilson liquid handler. ^1^H NMR spectra were recorded, with the DMSO signal dampened. The bulk water signal at 4.7 ppm was selectively inverted. Data were processed using Bruker Topspin 3.14 ^65^.

### Fragment screening using a T2 relaxation-edited ligand-observed NMR assay

Cocktails of four structurally distinct compounds were created using MNova Screen software to ensure there was no significant overlap of peaks in the region of interest (5.5 – 9.5 ppm). Fragments were screened at 1 mM each, with 4% v/v DMSO. Compounds were dispensed in duplicate using an ECHO acoustic dispenser (0.5 μL of each, 100 mM stock in DMSO). TNKS2 ARC4 (35 μM in 25 mM HEPES-NaOH pH 7.5, 100 mM NaCl, 2 mM TCEP, 10% D_2_O) was added to one cocktail, and buffer alone to the other replicate for a control sample of compounds alone. Mixtures were incubated for 20 min at room temperature, and then transferred into 1.7 mm NMR tubes. ^1^H relaxation-edited NMR spectra were collected with double solvent suppression applied to dampen the water and DMSO solvent signals. ^1^H spectra of each individual compound were used as reference spectra.

Data were processed using Bruker Topsin 3.14, then analysed using MNova Screen software. Only peaks between 5.5 and 9.5 ppm were considered. Peaks with a height of <5% maximum peak height within the region of interest were considered to be noise, and the minimum matched peak level was set at >51%. The relative peak intensity change (I) was calculated by equation 1 for all peaks in the 5.5 – 9.5 ppm region, for each compound.

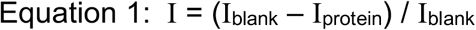

The average percentage change was then calculated, and compounds designated a hit if ≥26% reduction in signal (2 σ). Hit fragments were split into two groups: those with a signal change ≥39% (3 σ, 35 compounds), and those with a signal change of 26 – 39% reduction (2 – 3 σ, 65 compounds). Compounds of the first hit group (>3 σ) were tested individually in a second relaxation-edited assay, and the second hit group was tested in a waterLOGSY experiment, as this should rescue any genuine binders with a relatively small signal in the relaxation-edited assay, which has an intrinsic variability of ±10%. Protein ^1^H spectra of TNKS2 ARC4 (200 μM) were measured at 24 h intervals to ensure protein stability and folding for the duration of screening experiments.

### Mass spectrometry

#### Fragment quality control

An Agilent 6520 Quadrupole time of flight (qToF) mass spectrometer, with a 1200 series HPLC, fitted with an ESI/APCI multimode ionisation source was used. All solvents were modified with 0.1% formic acid. Fragments (2 mM in DMSO) were injected (2 μL) onto a Purospher STAR RP-18 end-capped column (3 μm, 30 × 4 mm, Merck KGaA). Chromatographic separation was carried out over a 4-min gradient elution (90:10 to 10:90 water:methanol) at 30 °C. UV-Vis spectra were measured at 254 nm on a 1200 series diode array detector (Agilent). The eluent flow was split, with 10% infused into the mass spectrometer. Eluent and nebulising gas were introduced perpendicular to the capillary axis, and applying 2 kV to the charging electrode generated a charged aerosol. The aerosol was dried by infrared emitters (200 °C) and drying gas (8 L/min of N_2_ at 300 °C, 40 psi), producing ions by ESI. Aerosol and ions were transferred to the APCI zone where solvent and analyte were vaporized. A current of 4 μA was applied, producing a corona discharge between the corona needle and APCI counter electrode, which produced ions by APCI. ESI and APCI ions simultaneously entered the transfer capillary along which a potential difference of 4 kV was applied. The fragmentor voltage was set at 180 V and skimmer at 60 V. Mass spectrometry data were acquired in positive ionisation mode over a scan range of m/z 160-950 with reference mass correction at m/z 622.02896 (Hexakis(2,2-difluoroethoxy)phosphazene). Data was analysed using MassHunter Qualitative Analysis B.06.00 (Agilent). Compound purity was calculated using the highest value of %UV (at 254 nm) or %TIC (total ion count).

### Protein expression and purification

Tankyrase ARC constructs were produced as previously described (see Pollock *et al*, 2017 for construct details) ^49^. Uniformly ^15^N-labelled protein was grown in M9 minimal media containing ^15^NH_4_Cl (CK isotopes). One litre of M9 minimal media was prepared by combining M9 medium (10X stock, 100 mL), trace elements solution (100X, 10 mL), glucose (20% w/v, 20 mL), magnesium sulfate (1 M, 1 mL), calcium chloride (1 M, 0.3 mL), biotin (1 mg/mL, 1 mL), thiamin (1 mg/mL, 1 mL) and making up to 1 L with water. M9 medium (10X) contained disodium hydrogen phosphate (60 g/L), potassium dihydrogen phosphate (30 g/L), sodium chloride (5 g/L), and ^15^N ammonium chloride (25 g/L).

BL21-CodonPlus (DE3)-RIL *E. coli* cells were transformed with a pETM30-2 plasmid containing the gene for a His_6_-GST tagged human tankyrase ARC construct ^4^. A single colony was selected and amplified in LB media (5 mL, Laboratory Support Services, ICR) for 6 h. This culture (1 mL) was then used to inoculate minimal media (200 mL) containing kanamycin (50 μg/mL) and chloramphenicol (34 μg/mL), and grown at 37 °C overnight. This starter culture (25 mL) was then used to inoculate each litre of minimal media, containing antibiotics as before. Cultures were grown at 37 °C with shaking (180 rpm) to an optical density of 0.6, measured at 600 nm. The temperature was reduced to 18 °C, and protein expression was induced by the addition of isopropyl-β-D-1-thiogalactopyranoside (IPTG) (0.5 mM). The cultures were incubated at 18 °C overnight. Cells were harvested by centrifugation (4000 × g, 30 min). The pellet was stored at −80 °C until purification following the previously described method ^49^.

Doubly ^15^N/^13^C-labeled protein for the backbone and partial side-chain assignment of TNKS2 ARC4 was prepared as the ^15^N-labeled protein, except that ^13^C-D-glucose (Cambridge Isotope Laboratories; at 6 g/L of M9 media) was used as well. Method details will be reported elsewhere (Zaleska *et al*., 2019) ^69^.

### K_d_ determination using chemical shift perturbation

#### TBM peptide and fragment titrations

^15^N-TNKS2 ARC4 (488-649) (300 μM final concentration; 5 μL of 3 mM stock) in NMR buffer (45 μL, 25 mM HEPES-NaOH pH 7.5, 100 mM NaCl, 1 mM TCEP, 10% D_2_O) was used as the baseline sample. Peptide titration samples were prepared by diluting peptide (table 5) with NMR buffer (45 μL), then adding TNKS2 ARC4 (300 μM final concentration; 5 μL of 3 mM stock). Separate samples were prepared for each concentration point.

**Table 5.** Peptide concentrations for titration in protein-observed NMR.

|  |  |  |  |  |  |  |
| --- | --- | --- | --- | --- | --- | --- |
| [peptide] (μM) | 50 | 150 | 300 | 600 | 900 | 1500 |
| ratio peptide:protein | 1:6 | 1:2 | 1:1 | 2:1 | 3:1 | 5:1 |

Fragments titration samples were prepared by diluting fragments in NMR buffer (25 mM HEPES-NaOH pH7.5, 100 mM NaCl, 1 mM TCEP, 10% D_2_O) and backfilling with DMSO to keep a constant DMSO concentration of 5% (table 6). ^15^N-TNKS2 ARC4 (300 μM final concentration; 5 μL of 3 mM stock) was added. Protein with 5% DMSO alone was used as the baseline. Separate samples were prepared for each concentration point. ^1^H-^15^N HSQC spectra were acquired over 3 h, with 64 scans and a spectrum width of 16.00 ppm for ^1^H and 29.00 ppm for ^15^N.

**Table 6.** Fragment concentrations for titration in protein-observed NMR.

|  |  |  |  |  |  |  |  |
| --- | --- | --- | --- | --- | --- | --- | --- |
| [fragment] (μM) | 75 | 150 | 300 | 600 | 1200 | 2400 | 4800 |
| ratio fragment:protein | 1:4 | 1:2 | 1:1 | 2:1 | 4:1 | 8:1 | 16:1 |

The pH of the peptide and fragment stocks (at 3 mM) was confirmed to exclude the possibility that peak shifts were due to changes in pH during the titration.

#### Analysis of protein-observed NMR data

Data were processed in Bruker Topspin 3.14 and analysed using CcpNmr Analysis software v2.4.2 ^66^.

To enable identification of peptide and fragment binding sites, a full backbone and partial side-chain assignment of ^15^N-^13^C-labelled TNKS2 ARC4 (488-649) was performed. Overall, out of 165 amino acids (construct + 3 N-terminal residues introduced by the cloning method), 164 residues were assigned and backbone amides were missing for only two non-proline residues. Assignment details and methods will be reported elsewhere (Zaleska *et al*., 2019) ^69^.

Peaks that shifted were picked manually in each spectrum. The chemical shifts for each peak were measured and exported into Microsoft Excel, where the change in chemical shift from baseline was calculated for hydrogen and nitrogen shifts. The average Euclidean distance shifted (d) was then calculated using Equation 2, weighting the different nuclei:

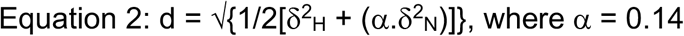

Values of d were plotted against ligand concentration in GraphPad Prism, and fitted with equation 3:

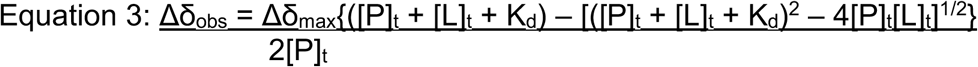

K_d_ values were calculated for each peak that shifted individually. The mean of all shifting peaks was then calculated to give an apparent K_d_ value ^67^.

#### K_d_ determination using isothermal titration calorimetry

An ITC200 (MicroCal) instrument was used, fitted with a twisted syringe needle, stirring at 750 rpm. All solutions were degassed using a ThermoVac before use. The reference cell was filled with buffer (200 μL, 25 mM HEPES-NaOH pH 7.5, 100 mM NaCl, 1 mM TCEP, 1% v/v DMSO). The cell was filled with TNKS2 ARC4 (488-649) (200 μL, 200 μM in identical buffer as above). 20 injections (1 × 0.5 μL, then 19 × 2 μL) of compound **9** (5 mM, 1% v/v DMSO in buffer) were performed, with 180 s between injections. Blank correction was performed by titrating compound into buffer alone (25 mM HEPES-NaOH pH 7.5, 100 mM NaCl, 1 mM TCEP, 1% v/v DMSO) using the same injection protocol as above. The first injections from each run were discarded from data analysis. Data were analyzed using Origin software with a one-site binding model. Titrations were repeated n=5. Global analysis was performed using SEDPHAT software ^68^.

#### *In-silico* prediction of fragment hotspots and pockets on TNKS2 ARC4

For the FTMap analysis, TNKS2 ARC4 chain D from the TNKS2 ARC4-3BP2 co-crystal structure (PDB 3TWR) ^4^ was submitted to the FTMap web server (fTMap.bu.edu) and analyzed under protein-protein interaction mode, as detailed under the published conditions ^59,60^.

For pocket identification using the Roll algorithm, TNKS2 ARC4 chain D from the TNKS2 ARC4-3BP2 co-crystal structure was submitted to the Pocasa 1.1 web server (altair.sci.hokudai.ac.jp/g6/service/pocasa/) and analyzed with the following parameters: probe radius, 2 Å; single point flag, 16; protein depth flag, 18; grid size, 1 Å; atom type, protein.

## Supporting information

Supplementary experimental information and four supplementary figures are available online.

### Acknowledgements

We thank Fiona Jeganathan for her help in optimizing the DSF assay, Meirion Richards for mass spectrometry analysis, and Rob van Montfort and Rosemary Burke for access to technologies. Work in the SG laboratory is supported by The Institute of Cancer Research (ICR), by Cancer Research UK through a Career Establishment Award (C47521/A16217), and by the Lister Institute of Preventive Medicine. Work in the IC laboratory is supported by ICR and by Cancer Research UK through funding to the Cancer Therapeutics Unit (C309/A11566). KP was supported by a Wellcome Trust PhD studentship (WT102360/Z/13/Z). This project was further supported by a Faringdon Proof of Concept Fund award from ICR.

## Author Contributions

Designed and planned experiments: KP, SG and IC; protein production: KP; fragment screening by thermal shift: KP; fragment screening and validation by NMR: KP and ML; fragment binding site location by protein-observed NMR: KP, MZ and MP; data analysis: KP, MZ, SG and IC; wrote the paper: KP, SG and IC with input from all authors.

