## Supplementary Information for "Fragment-based screening identifies molecules targeting the substrate-binding ankyrin repeat domains of tankyrase"

#### Supplementary Figure Legends

**Supplementary Figure 1.** Binding of TBM peptides containing arginine substituents, measured by fluorescence polarization (FP). Data shown are the mean of 2 independent experiments ± SD. Values were normalized with 0% and 100% set as the minimum and maximum values of the 3BP2 positive control. Data were fitted to a four-parameter variable-slope dose-response curve using GraphPad Prism 6.0. The insets show docking poses for R'-SP tripeptides returned by GOLD, with R substituent tripeptides shown in green and the original RSP tripeptide (from the 3BP2 TBM, 3TWR) shown in purple, both colored by heteroatom.

**Supplementary Figure 2. (A)** The effect of DMSO (0 – 10%) on the melting temperature ($T_m$) of TNKS2 ARC5 was determined by DSF. DMSO up to 10% v/v had no discernable effect on the $T_m$ of TNKS2 ARC5. **(B)** The correlation between $IC_{50}$ values measured by fluorescence polarization and melting temperature stabilization ($\Delta T_m$) as measured by DSF for the arginine substituent peptides. Non-binding peptides are shown in red, and peptides with a measurable $IC_{50}$ are shown in blue.

**Supplementary Figure 3.** $^1$H-$^{15}$N HSQC NMR spectrum, showing the chemical shift perturbations (CSPs) upon titration of the TBM peptide from 3BP2 at the indicated concentrations. Residues constituting the TBM binding site are color-coded as in Figure 1B. The concentration of TNKS2 ARC4 was 50 μM. Small panels show example CSPs displaying slow-exchange and fast-exchange kinetics.

**Supplementary Figure 4. (A)** $^1$H-$^{15}$N HSQC NMR spectrum, showing the CSPs upon compound **3** titration at the indicated concentrations. The concentration of TNKS2 ARC4 was 50 μM. **(B)** Same as (A) but for compound **5**. **(C)** Plot of CSPs in TNKS2 ARC4 induced by the addition of 16-fold excess of either compounds **3** or **5** ([TNKS2 ARC4] = 50 μM; [compound **3**/**5**] = 800 μM) or compound **9** ([TNKS2 ARC4] = 300 μM; [compound **9**] = 4800 μM). The residue labels (color-coded by TBM binding determinant: "central patch" or "aromatic glycine sandwich") correspond to CSPs of >2σ (>0.042) of the CSP average observed for compound **5**.

#### Supplementary Materials and Methods

##### Modelling of arginine replacements

ROCS software (version 3.2.0.4, OpenEye Scientific Software, Santa FE, NM, USA) [1] was used to screen a library of potential arginine replacements against the structure of arginine, looking for shape and pharmacophore matches. GOLD software [2] was used to dock the library of R'SP fragments with chain A (ARC4) from the TNKS2 ARC4 crystal structure in complex with 3BP2 peptide (PDB code 3TWR) [3]. Chain E (the corresponding 3BP2 peptide) was used as a template.

The TNKS2 ARC4 protein was prepared in MOE (Molecular Operating Environment, 2013.08, Chemical Computing Group Inc., Montreal, QC, Canada), and then uploaded into GOLD. The orientation of Asn/Gln/His side chains were as assigned in the crystal structure. The binding site was defined from coordinates at the centre of the arginine cradle. All residues within 10 Å were defined as the binding site. The crystal structure of the RSP extracted ligand was used as a reference ligand. The default scoring function settings were used. Protein rotatable bonds were fixed, and the proline at position three of the TBM was used as a scaffold to anchor the peptide. Twenty genetic algorithm runs were performed. The overall shape of the reference ligand (RSP from the crystal structure) was used as a similarity template for comparison of docked poses. Returned poses were exported into MOE and analysed manually by comparison with the crystal structure to rank peptides predicted to bind.

##### Synthesis of 3BP2 and arginine substituent peptides

N-Fmoc protected amino acids and Wang resin were purchased from Novabiochem. HATU was purchased from Apollo Scientific. Imidazole acids were purchased from Akos. All other reagents and solvents were acquired from Sigma Aldrich unless specified otherwise. An Activotec P14 peptide synthesiser, fitted with a 150 mL reaction vessel, was used.

The 3BP2 TBM peptide and arginine substituent peptides were synthesised using solid phase peptide synthesis (SPPS). A standard Fmoc SPPS procedure was followed, starting from Fmoc-Ser(tBu)-Wang resin. At each stage, 10 mL of solvent or reagent mixture was used per gram of resin, unless specified otherwise. The resin was swollen in *N, N*-dimethylformamide (DMF) for 15 min, before the 9-fluorenylmethyloxycarbonyl (Fmoc) protecting group was removed from the N-terminal amine. Piperidine (20% v/v in 1-Methyl-2-pyrrolidinone (NMP)) was added to the resin and agitated for 15 min. The resin was washed with DMF, and then the deprotection process was repeated. The resin was washed sequentially (5 min for each solvent) with DMF (x2), dichloromethane (DCM) (x2), methanol (MeOH) (x2), DCM, MeOH, DCM and diethyl ether.

The activated amino acid was prepared in a separate flask. Fmoc-Aaa(PG)-OH (4 eq.) was dissolved in DMF with 5% *N,N*-Diisopropylethylamine (DIPEA) (20 mL). 1-[Bis(dimethylamino)methylene]-1*H*-1,2,3-triazolo[4,5-*b*]pyridinium-3-oxid hexafluorophosphate (HATU) (3.6 eq.) was added and the mixture stirred at room temperature for 30 min. The activated amino acid was added to the resin and shaken gently for 2 h. The resin was washed with DMF (x2), and then the amino acid activation and coupling steps were repeated. The resin was washed sequentially as above. This process was repeated to extend the peptide chain to the desired length.

TFA:triethyl silane (TES) 95:5, (1 mL / 50 mg resin) was added to the resin and shaken for 2 h. The TFA mixture was then filtered drop-wise into diethyl ether (20 mL) cooled to -20 °C, which induced peptide precipitation. The precipitate was pelleted by centrifugation (13,300 rpm, 5 min). The solution was decanted and the precipitate dried under vacuum, to yield the solid peptide. Peptide identity was confirmed by mass spectrometry analysis, and then peptides were dialysed into buffer (25 mM HEPES-NaOH pH 7.5, 100 mM NaCl, 2 mM TCEP). Amino acid analysis by the PNAC facility at Cambridge University gave an accurate concentration of each amino and therefore the peptide, and further confirmed peptide identity and purity. Peptide samples were flash frozen in liquid nitrogen and stored at -80°C before use.

**Analysis of peptides by mass spectrometry (MS)**

An Agilent 6210 Time of flight (ToF) liquid chromatography mass spectrometer (LC-MS) was used for analysis of peptides up to 800 Da. A gradient of 90% aqueous + 1% formic acid (solvent A) to 90% methanol (solvent B) was run over 15 min. Peptides with molecular weight larger than 800 Da were analysed using an Agilent 6520 Quadrupole time of flight (qToF) LC-MS to determine the accurate mass. Both instruments had a 1200 series LC. A dual ESI ionisation source, and a Jupiter column (25 x 0.46 cm, C18, 5 μm pore size, Phenomenex) were used. A gradient of 90% aqueous + 1% formic acid (solvent A) to 90% acetonitrile + 1% formic acid (solvent B) was run over 15 min. A blank sample of water was run on the column prior to any samples being run. Data was analysed using MassHunter Qualitative Analysis (Agilent).

**Fluorescence polarization assay**

Unlabeled arginine substituent peptides were titrated up to 1 mM with 5 nM probe (Cy5-RSPPDGQS, JPT Peptide Technologies) and 5 μM TNKS ARC4. 3BP2 16-mer was used as a positive control, and the same peptide with the glycine at position 6 mutated to arginine (G6R) was used as a negative control. Plus F ProxiPlates (PerkinElmer) were used with a 2103 Envision Multilabel plate reader (PerkinElmer), fitted with filters 620 nm (excitation) and 688 nm (emission). Filter bandwidths were 10 nm. A Nanodrop 2000 was used for measuring protein and probe concentrations by UV absorption. TNKS2 ARC4 protein thawed from -80 °C, and centrifuged at 13,000 x *g* before use.

Assays were performed in technical duplicate. A 2× stock solution containing TNKS2 ARC4 protein (10 μM) and fluorescently tagged probe (10 nM) in assay buffer (25 mM HEPES-NaOH pH 7.5, 100 mM NaCl, 1 mM TCEP, 0.01% CHAPS) was prepared. A 2-fold dilution series of untagged peptide in assay buffer was made (0 – 2 mM). Protein and peptide stock solutions were mixed 1:1. Blank samples contained assay buffer only. The plate was sealed, centrifuged for 1 min at 1000 × *g*, and then incubated in the dark for 30 min before reading. Readings were taken with 50 flashes per well, and measured in millipolarisation (mP) units. Data was exported to Microsoft Excel, and analysed in GraphPad Prism 6.0. The mean of technical duplicate values was calculated, and FP values were plotted against unlabeled peptide concentration. Curves were normalized using the maximum and minimum FP values of the control 3BP2 TBM peptide, before data from independent experiments were combined. Curves were fitted using a non-linear regression, log(inhibitor) vs. response – variable slope (four parameters) model.

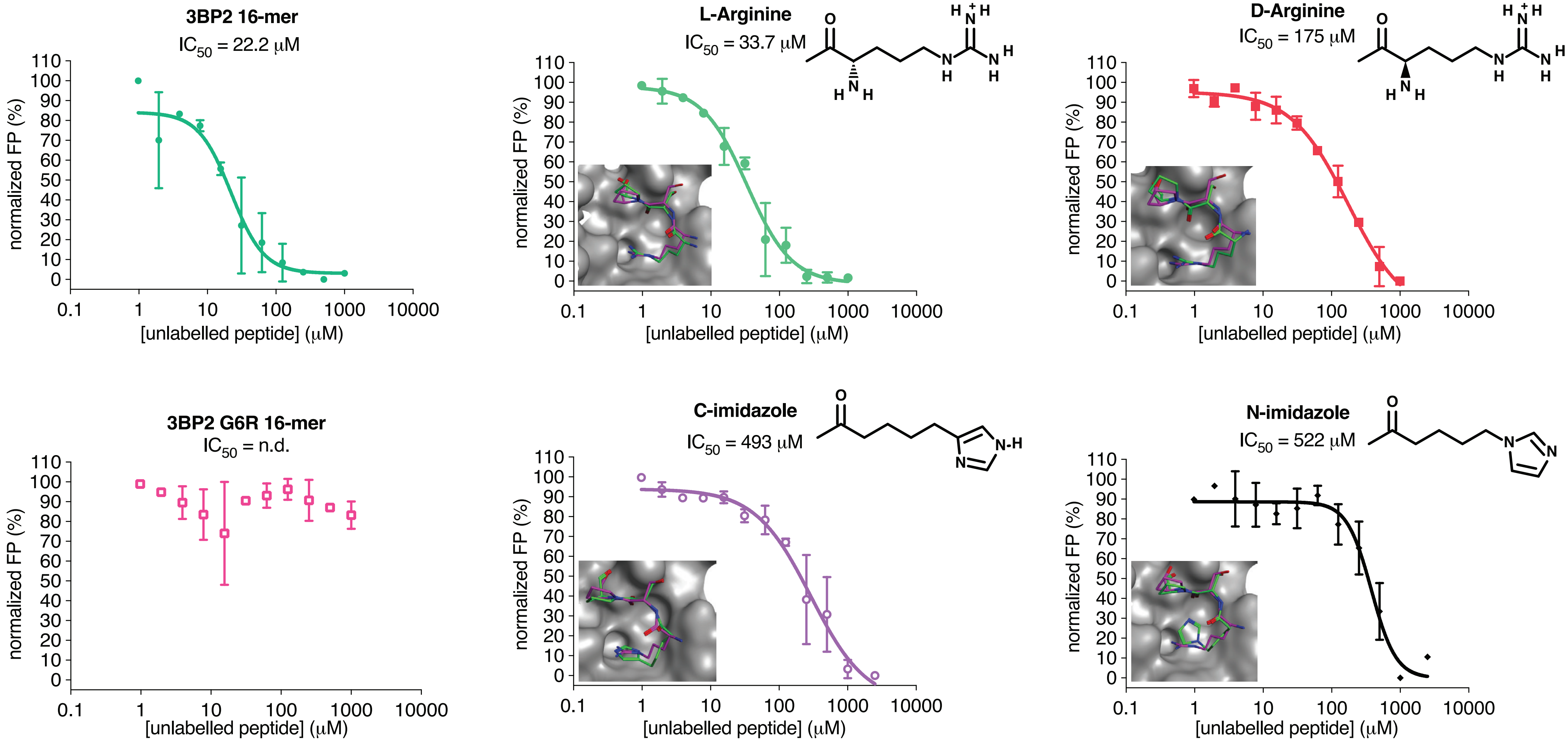


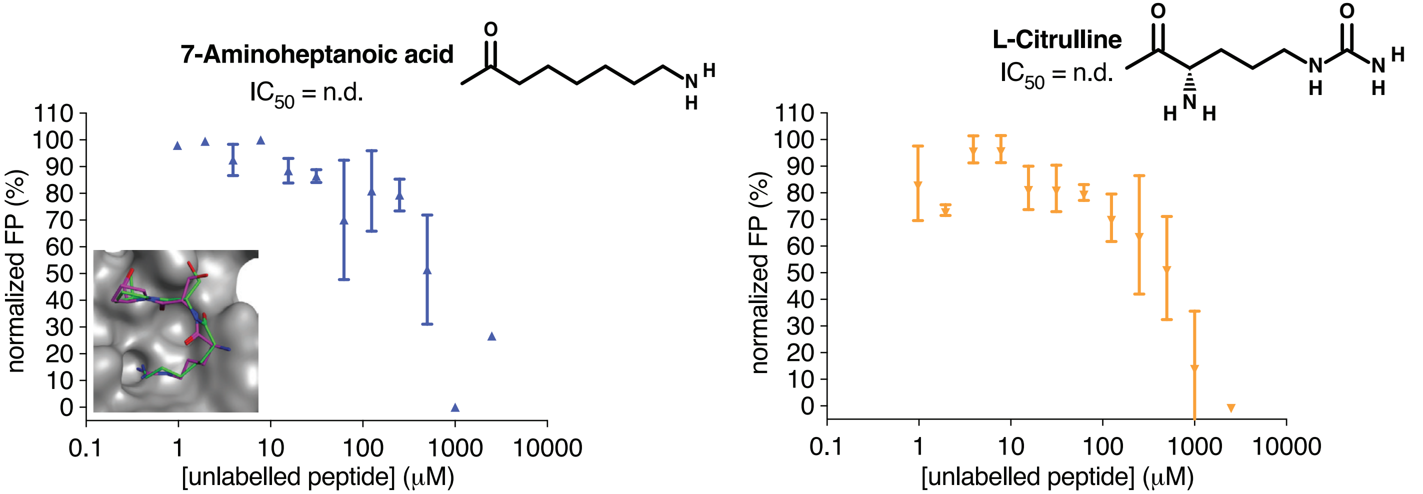


**Supplementary Figure 1**

**A**

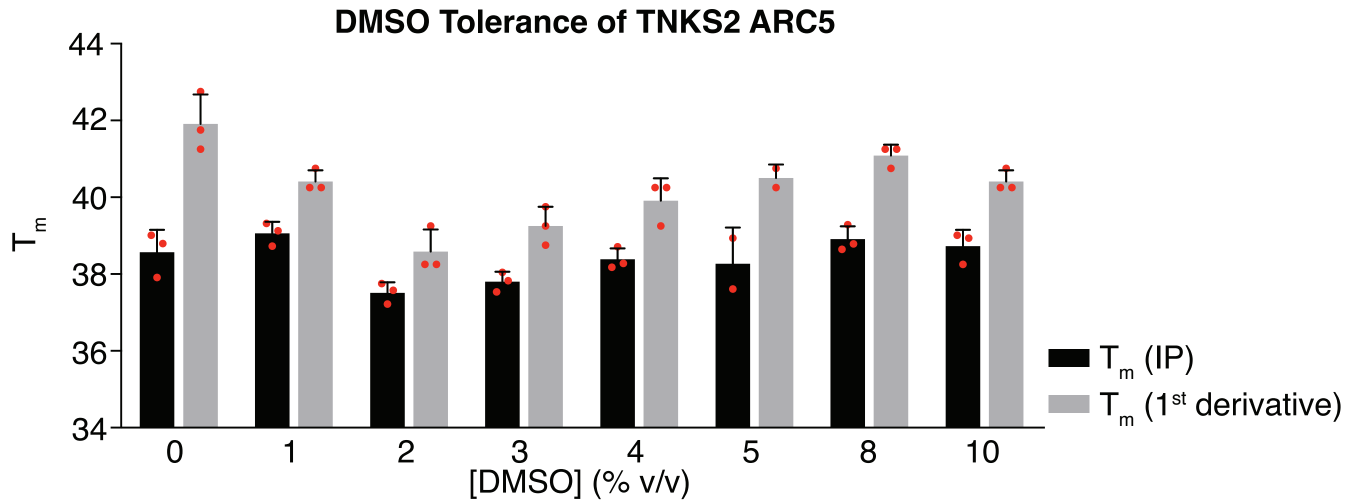


**B**

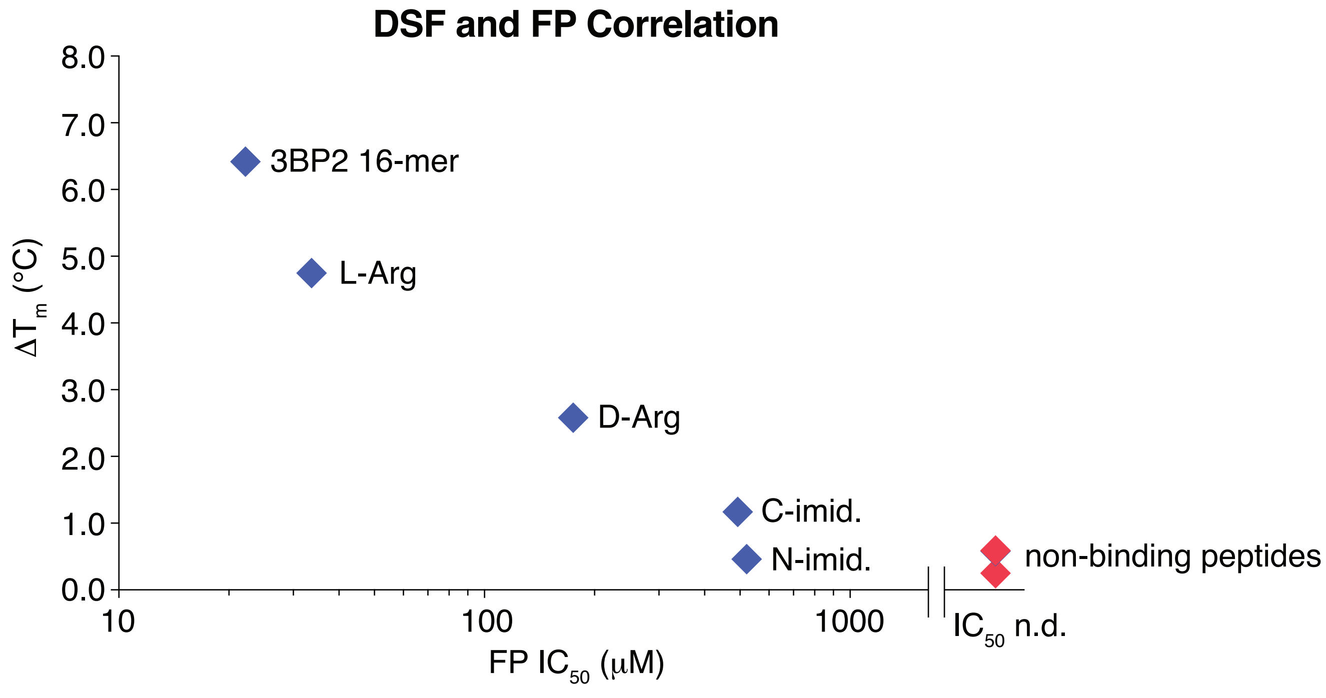


### Supplementary Figure 2

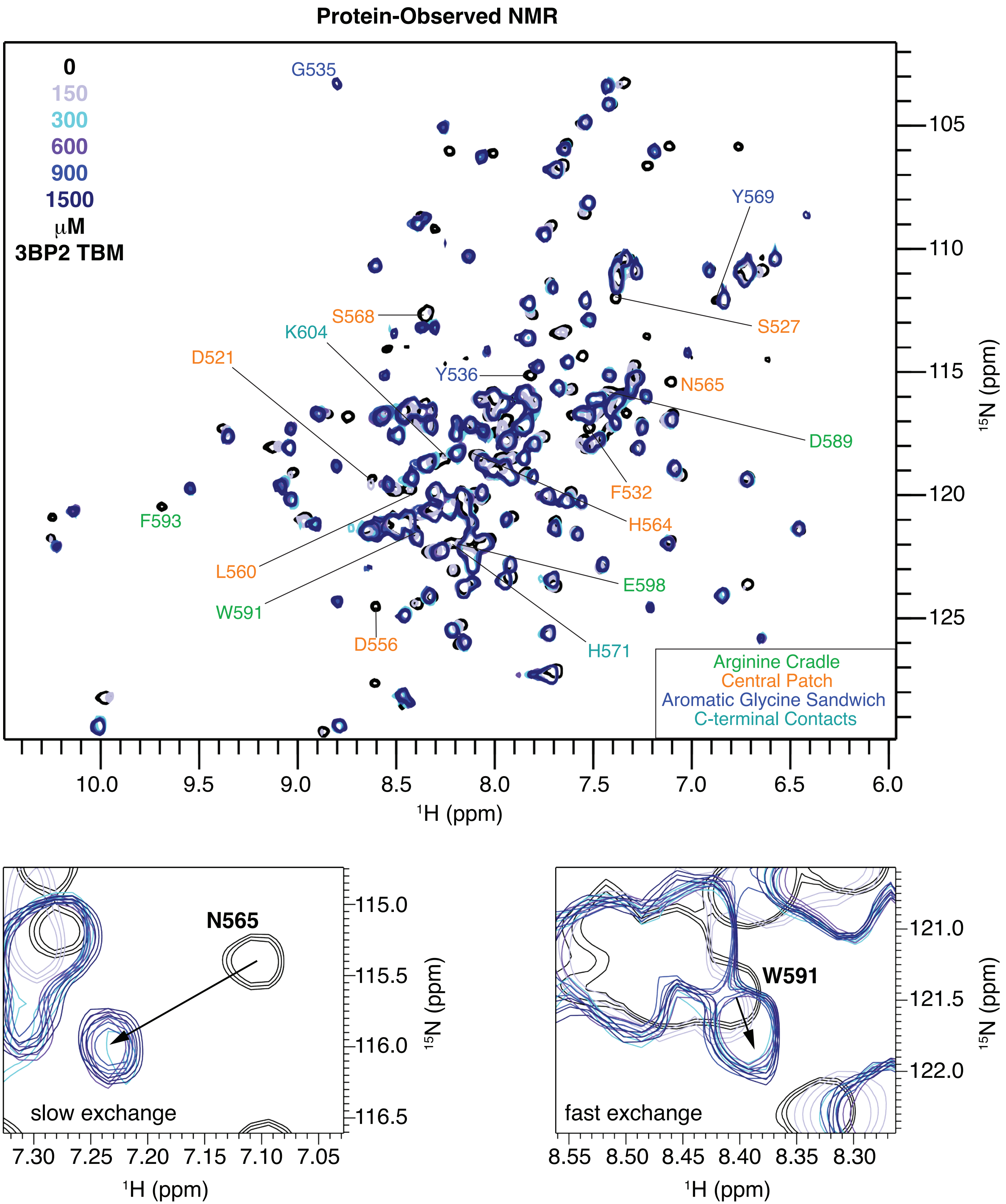


**Supplementary Figure 3**

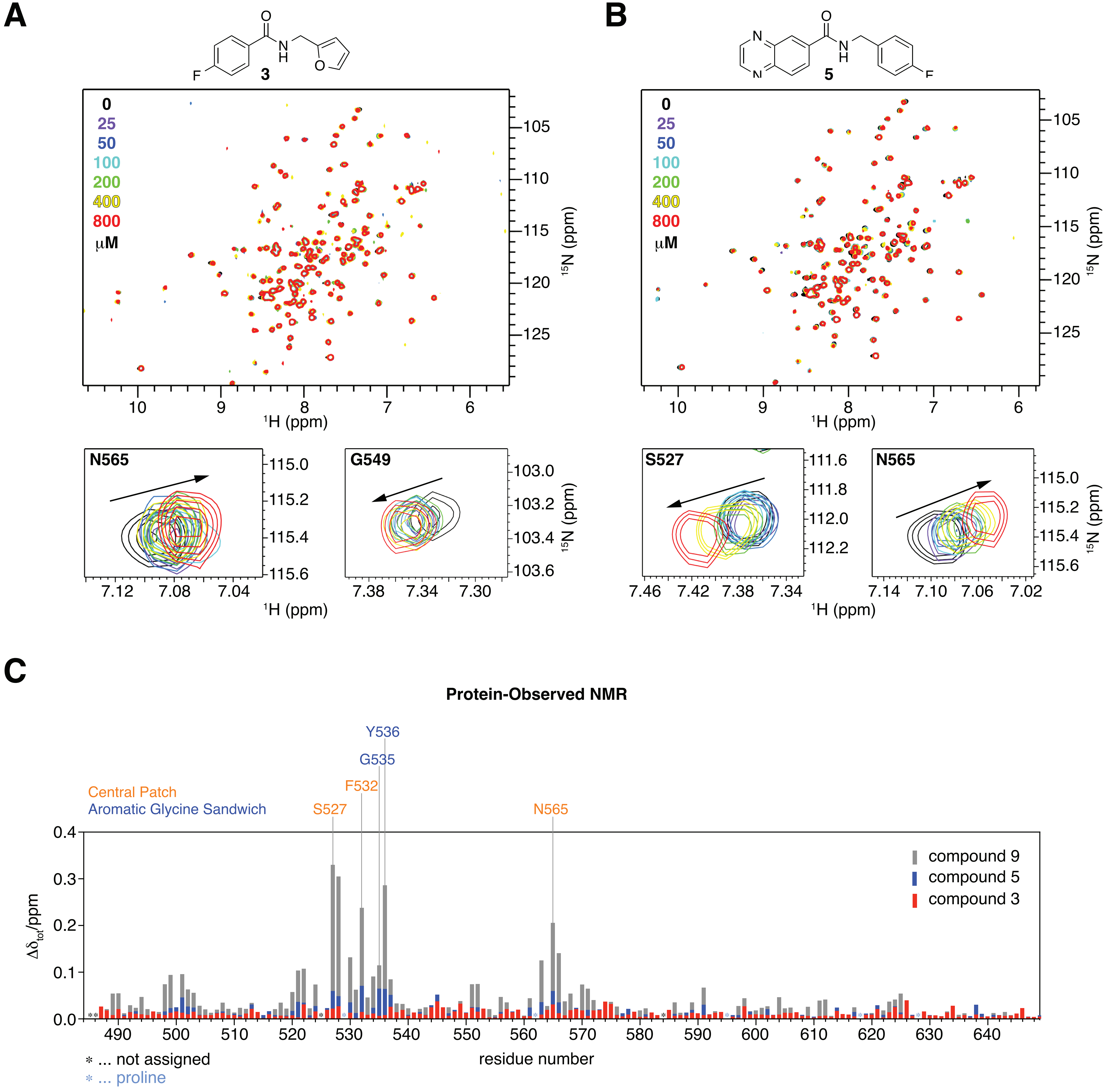


**Supplementary Figure 4**
